# Great Expectations: A Critical Review of and Recommendations for the study of Reward Processing as a Cause and Predictor of Depression

**DOI:** 10.1101/2020.03.04.975136

**Authors:** Dylan M. Nielson, Hanna Keren, Georgia O’Callaghan, Sarah M. Jackson, Ioanna Douka, Charles Y. Zheng, Pablo Vidal-Ribas, Narun Pornpattananangkul, Christopher C. Camp, Lisa S. Gorham, Christine Wei, Stuart Kirwan, Argyris Stringaris

## Abstract

Both human and animal studies support the relationship between depression and reward processing abnormalities, giving rise to the expectation that neural signals of these processes may serve as biomarkers or mechanistic treatment targets. Given the great promise of this research line, we scrutinize those findings and the theoretical claims that underlie them. To achieve this, we apply the framework provided by classical work on causality as well as contemporary approaches to prediction. We identify a number of conceptual, practical, and analytical challenges to this line of research, and use a pre-registered meta-analysis to quantify the longitudinal associations between reward processing aberrations and depression. We also investigate the impact of measurement error on reported data. We find that reward processing abnormalities do not reach levels that would be useful for clinical prediction, yet the evidence thus far does not exclude their possible causal role in depression.

## Introduction

Aberrations in how people form expectations about reward and how they respond to receiving rewards are thought to underlie depression, in particular the symptom of anhedonia. Anhedonia is the loss of interest or pleasure in activities that were previously considered pleasurable (1,2). Several lines of evidence support the relationship between reward and anhedonia. The most basic of these is face validity between anhedonia and reward related processes. These processes are instantiated in a network encompassing the ventral striatum, the anterior cingulate cortex (ACC), and the orbital prefrontal cortex (OFC) (3) and work from animal models has shown that lesions in these areas produce anhedonic phenotypes (4,5). Meta-analytic evidence from fMRI and EEG studies concurs; reduced neural signals in these brain areas acquired during reward tasks are associated with depression (6–8).

These findings offer hope for the use of reward processing aberrations as either biomarkers for prediction of depression onset and course, or as targets for future treatments. Given their potential, we scrutinize those findings and the theoretical claims that underlie them. To achieve this, we build on previous reviews that have quantified cross-sectional associations; evaluating the literature in the framework provided by classical work on causality (9) and contemporary approaches to prediction (10). First, we critically examine the conceptual challenges to the purported relationship between reward processing and depression. Second, we consider the challenges of measuring symptoms of depression and reward processing. Third, we examine the existing meta-analytic evidence for a cross-sectional association between reward processing and depression. Fourth, we present a new meta-analytic analysis of the critical longitudinal associations and estimate the magnitude of these effects in the best case. Fifth, we review any evidence on the impact that manipulation of reward processes has on depression. Finally, based on the challenges that we have identified, we provide a list of specific recommendations about how to improve modelling and experimental approaches to the study of the relationship between neural signals of reward processing and depression.

## Conceptual challenges

Reward processing encompasses a number of phases that we illustrate using the following example: A child sees a chocolate wrapper on her kitchen table and forms the *expectation* that there is chocolate nearby. In other words, she forms a *prediction* and *anticipates* that she will find chocolate. The terms expectation, prediction and anticipation are often used interchangeably; here we will largely refer to prediction, but many tasks that probe this behavior refer to it as the anticipatory phase. The child *decides* to investigate a nearby tin on the kitchen table and then tries to open it based on her prediction that it may contain chocolate. Here, the child *acts* and expends *effort* to obtain a reward. The child finds one chocolate left and enjoys the *experience* of eating the chocolate. The next day, she finds a new tin of chocolates, but opens it to discover that all of the chocolates have been eaten by her brother and becomes disappointed. The child experiences what is termed a negative *reward prediction error (RPE)*, in this case a *negative prediction error*, because she got less than expected. RPEs are thought to underlie reward-related learning, which for this example, would reduce the likelihood that the child looks into the tin for chocolate in the future.

The disruption of any of these reward processes - prediction, decision, effort, and experience - are thought to be associated with symptoms of anhedonia (7). Indeed, it is argued that anhedonia in the context of depression may be of the same kind as anhedonia experienced when physically ill (11,12).

Motivated by this thinking--and by preclinical evidence from animals and humans--there has been a surge of experimental research in humans that uses reward tools to understand depression. Here, we critically review some of the conceptual challenges to this research.

### Theoretical weakness

As we attempt to probe the causal nature of the relationship, specific theories for the mechanism by which reward processing causes depression should be postulated and tested. For example, we hypothesize that reduced neural reward processing signal at time t predicts future changes in anhedonic symptoms only through the persistence of that reduction (Figure 1, Hypothesis). When we test this hypothesis, failure to falsify it does not confirm the model since there are alternatives that may also be supported by the data (Figure 1, Alternatives):

**Figure 1:**
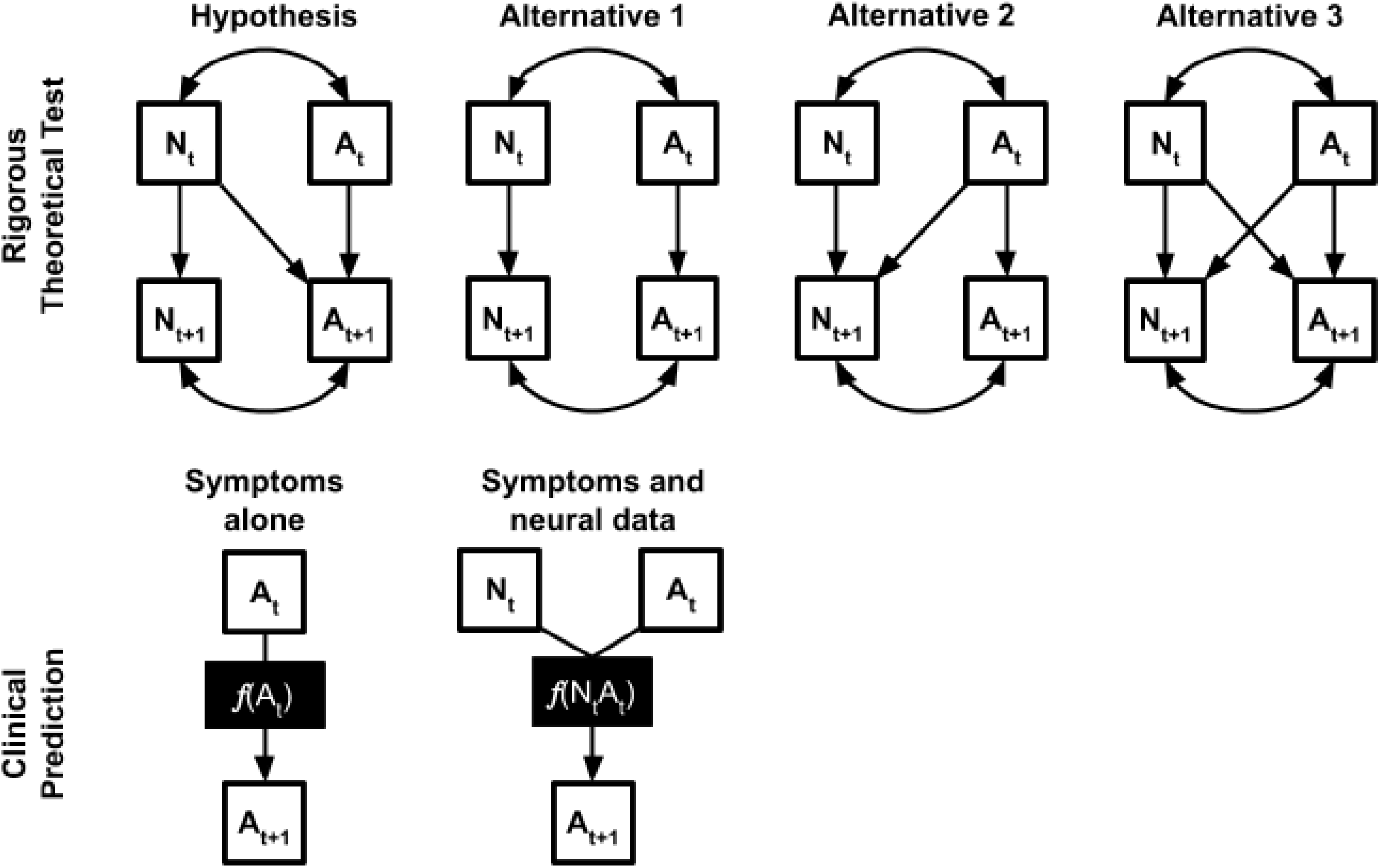
Two approaches for testing the relationship between neural signals of reward processing (N) and anhedonic symptoms (A). The first is an example of a rigorous theoretical test of our hypothesis that reduced neural reward processing signal at time t predicts future changes in anhedonic symptoms only through the persistence of that reduction. We test that model for non-equivalence against three reasonable alternatives: 1) Two processes correlated cross-sectionally with no prediction across time. 2) The relationship is the opposite of our hypothesis, anhedonic symptoms predict reward processing signal. 3) A fully cross-lagged model where symptoms and reward processing interact over time. If we find our hypothesis to be superior, then we have strong evidence in its support. Alternatively, we could focus on making clinical predictions without concern for theoretical justification (note this is not an SEM). In this case, we are just trying to use baseline neural data (N_t_) to improve our prediction of future anhedonia (A_t+1_) beyond what we could already predict based on baseline anhedonia (A_t_) using some machine learning approach such as a regularized regression.

1. Two processes correlated cross-sectionally with no prediction across time.
2. The relationship is the opposite of our hypothesis, anhedonic symptoms predict reward processing signal.
3. A fully cross-lagged model where symptoms and reward processing interact over time.

If we showed that our *a priori* hypothesis was superior to reasonable alternatives, we would have stronger evidence to confirm our model (13). Alternatively, we could focus purely on clinical application without concern for theory; for example, testing if baseline neural reward signals improve the prediction of future anhedonia beyond what baseline anhedonia contributes (Figure 1, Clinical Prediction) -- a concept termed incremental validity (14). In both cases stronger evidence is provided by pre-registered hypotheses and analytical plans.

### State vs trait

A rarely discussed question is whether reward processing abnormalities are trait or state problems.. Longitudinal studies (reviewed in detail below) suggest that neural signal reduction precedes the onset of depression in some youth, which would suggest that it is a trait rather than a state condition. In addition, evidence of blunted reward processing in remitted MDD (15–17) and in unaffected first degree relatives (18) argue also for a trait interpretation. However, treatment studies find that aberrant neural reward processing normalizes over the course of treatment in correlation with symptom reduction (19,20), indicating that reward processing abnormalities may be a state rather than a trait. To our knowledge, no studies have directly tested how reward processing abnormalities covary with changes in anhedonia over the course of multiple assessments. If these are trait abnormalities, then their utility as biomarkers of the course of mental illness may be limited.

### Developmental moderation

There are good reasons to assume that development could somehow influence the association between reward and depression. For example, a dramatic rise in depression cases occurs during puberty (21) coinciding with a period of time when, normatively, adolescents are apparently more sensitive to rewards (22). Moreover, there is some meta-analytic evidence (7) to suggest that reduced reward processing in depression may be more pronounced in adolescents compared to adults. Yet, with notable exceptions (23,24), rarely are specific theories being proposed about the interplay of development with reward processing and depression (23,24). It is even rarer to see any robust tests of such theories (23). This is not surprising given how difficult it is to conduct such studies. For example, a study about the effects of puberty would ideally have the following characteristics: a) be large enough to test for interactions between developmental time (linear and quadratic) with reward processing for the outcome of depression; b) include puberty-specific measures (e.g. pubertal hormones) as well as environmental factors (e.g. school transition, bullying, etc) that could confound such relationships; and c) be representative of the population of adolescents. To our knowledge, the study that most closely fits these criteria is the Adolescent Brain and Cognitive Development (ABCD) study (25). We urge researchers to take advantage of the measures of development that will be available in the next rounds of data released by ABCD.

### Specificity

Often implicitly, but also explicitly (e.g. (11)), anhedonia in depression is thought of as an overall reduced wanting or liking of rewards. However, this model is challenged by the well-validated clinical observation that participants with depression have a higher likelihood of drug and alcohol addictions to hedonic substances (26). This suggests that reward processing abnormalities may present context-dependent dysregulation of hedonic processing as suggested by Volkow (27). This is a more nuanced and complex hypothesis to specify and is rarely, if ever, empirically tested.

Additionally, it is unclear whether reward processing abnormalities--for example, a reduction in the neural signal during the anticipation of a reward--are confined to anhedonia in the context of depression. There are several alternative hypotheses that have only partially been tested. First, within depression it remains to be established whether reward processing abnormalities are differentially related to anhedonia as opposed to other symptoms. We only know of two studies, which have found that anhedonia but not low mood are related to reward processing abnormalities in community (not clinically diagnosed) samples (28,29). Yet comparing anhedonia to other plausible symptoms, such as loss of energy or fatigue, is also important. Second, anhedonia is present in other disorders, such as schizophrenia or ADHD. Indeed, reduced striatal BOLD signal during reward anticipation has been described in these populations (30,31). In some studies, this signal has been accounted for by depression comorbidity (32); in others, this reduction was only observed in adult but not youth samples (33). In a recent study from our group, reduction in striatal activity was observed only in children with anhedonia but not in those with anxiety or ADHD in a community sample (whilst ADHD was associated with BOLD signal aberrations during a working memory task) (29).

## Measurement Challenges

### Measurement of the clinical phenotype

Human self-report, on which studies rely, is extremely important and it would be wrong--from a clinical as well as ethical point of view--to dismiss it. However, it is also fraught with the following problems, among others (see Rizvi *et al.* (2) for a thorough review of challenges).

#### Retrospective accounts of anhedonia

There are inherent problems with self-report of anhedonia, in particular consummatory anhedonia, or the lack of enjoyment when experiencing a reward. In the example above, the child would be asked--now sitting in some research laboratory--about her experience of consuming the chocolate. This requires forming the mental representation of a past event and attaching value to it, a different process than that of actual consummation, and in some ways more related to the process of predicting the value of a future reward based on past experiences. This is especially problematic in depressed patients, who tend to recall rewards as rarer than they actually were (34). Ecological momentary assessments may allow more direct measurement of consummatory anhedonia (35,36).

#### Human-animal translation

Self-report measures of anhedonia are obviously not translatable to animal models, so animal research into anhedonic symptoms focuses instead on behavioral assessment. However, these animal behavioral assessments don’t translate back to humans. A classical test of consummatory anhedonia in animals is the sucrose preference test--chronic stress reduces an animal’s preference for sucrose, which has been shown to reverse with the administration of antidepressant treatment. Yet, no differences between depressed and non-depressed participants have been found in a human sucrose test (37) and anhedonia is associated with failure to respond to SSRIs in human depression (38). This has cast doubt on whether reward consummation is a part of reward processing that is affected in depression. On the other hand, prediction has been assessed with analogous modalities in both animals and humans and was shown to be affected by depression, but the relationship with deficits of anticipation has not been demonstrated (39,40).

### Measurement of reward processing

Reward processing is a promising avenue for understanding the mechanisms of anhedonia, and several experimental approaches have been developed to isolate components such as anticipation or consummation of reward. Many behavioral tasks correlate poorly with self-report measures due to low reliability and measurement of different entities (41), and task-based functional magnetic resonance imaging (fMRI) may have similarly low reliability (42). Measurement of reward processing faces these problems and others:

#### Not measuring behavior

Some of the most widely used tasks in reward processing neuroimaging lack a behavioral output. For example, the titrated Monetary Incentive Delay (MID) task (43,44), an experimental set up used by us and many others in the field, does not offer behavioral output that could help differentiate between depressed and non-depressed participants. Interpreting blood oxygen level dependent (BOLD) signal in its own right is fraught with ambiguities: a reduced BOLD signal could be a deficit or a compensatory mechanism, and in the absence of a task-related behavioral response it becomes harder to interpret.

#### Measuring some but not all phases of reward processing

Most studies employ tasks that only measure some of the components of reward processing. For example, in the MID, the most commonly used task, only prediction (measured as neural activity during the anticipatory period) and experience (neural activity during the feedback period) of reward are probed, while other important phases such as decision and effort are left out (45). This means that key components of the reward system are not probed in the same individuals, and therefore inferences drawn about reward processing may be biased or partial. Computational modeling (as in (46,47)) of all of the phases of reward, potentially across multiple tasks within the same individuals, would allow a more thorough phenotyping of the reward system (48,49).

#### Multiplicity of measurement

Different neuroimaging studies define the same phase of reward processing in different ways. For example, the label *reward anticipation* is applied to analyses that contrast it to a neutral condition, a loss condition, or even just baseline activity. Taking just the fMRI studies reviewed in Keren, O’Callaghan *et al.* (7) and Ng *et al.* (6) as examples, we found 22 different tasks, half of which have only been used once (Figure 2, Table S1). Across these tasks, at least 64 different task-contrast combinations were used, 50 of them only once. The most commonly reported was the gain anticipation versus neutral anticipation contrast for the MID task in 10 studies. Given such a large space of potential tasks, contrasts, and analytical approaches, it is impossible to know if the contrasts and analyses used in any given paper are the only analyses done or if they are the result of searching that space for a significant finding. The rationale for these choices is often not stated, and when it is mentioned, there is no way of knowing if that justification is post-hoc.

**Figure 2:**
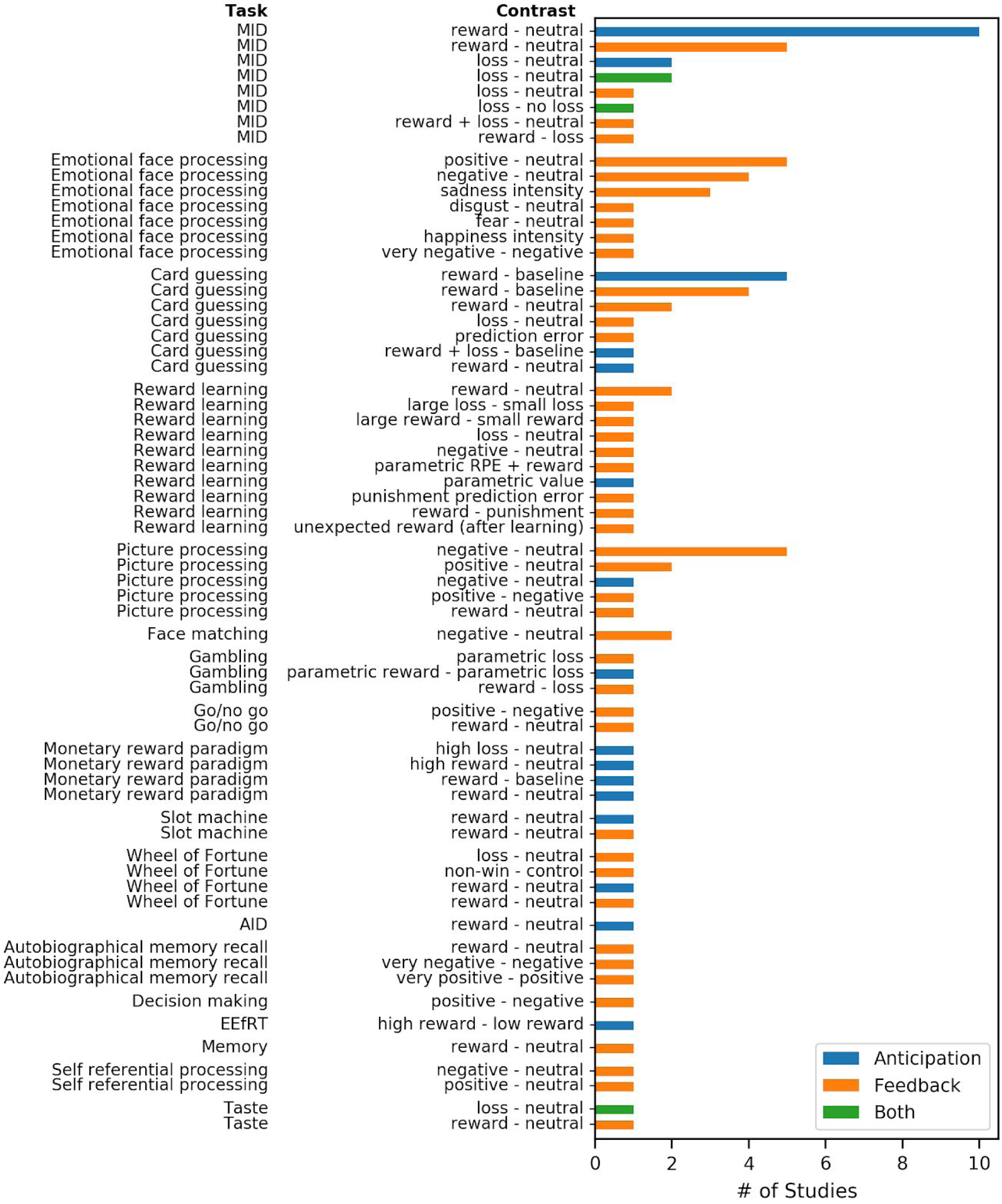
Diversity of tasks and contrasts in studies reviewed by Keren, O’Callaghan *et al.* (7) and Ng *et al.* (6). The majority of task/contrast combinations in the literature have appeared in only a single study. Classification of the task as assessing anticipation or feedback is based on the information reported in each meta-analysis. MID: Monetary Incentive Delay task; AID: Affect Incentive Delay; EEfRT: Effort-Expenditure for Rewards Task; RPE: Reward Prediction Error

## Cross-Sectional Association

Association may not imply causation but, equally, two variables cannot be causally related unless they correlate with each other. Moreover, a predictor variable that is strongly associated with a clinical outcome is more likely to be suited for use as a diagnostic or prognostic tool. In this section, we critically review work that has summarized the association between depression and reward processing and evaluate the clinical relevance of that association.

### fMRI

Three meta-analyses have examined the association between depression and reward processing aberrations as measured using fMRI during a reward-related task, such as the MID. All three papers focused on cross-sectional differences in reward processing between healthy volunteers and individuals with depression or participants at high risk of depression and they identified reduced response to reward in the ventral striatum or caudate during reward prediction or experience (6–8). The results are encouraging in that they are attained with the inclusion of different studies (Figure 3). Yet, a major disadvantage of the activation likelihood estimation (ALE) approach that was used in these studies is that it does not provide an estimate of the strength of association (50) and that studies with null effects cannot be included, introducing the possibility of a positive bias.

**Figure 3:**
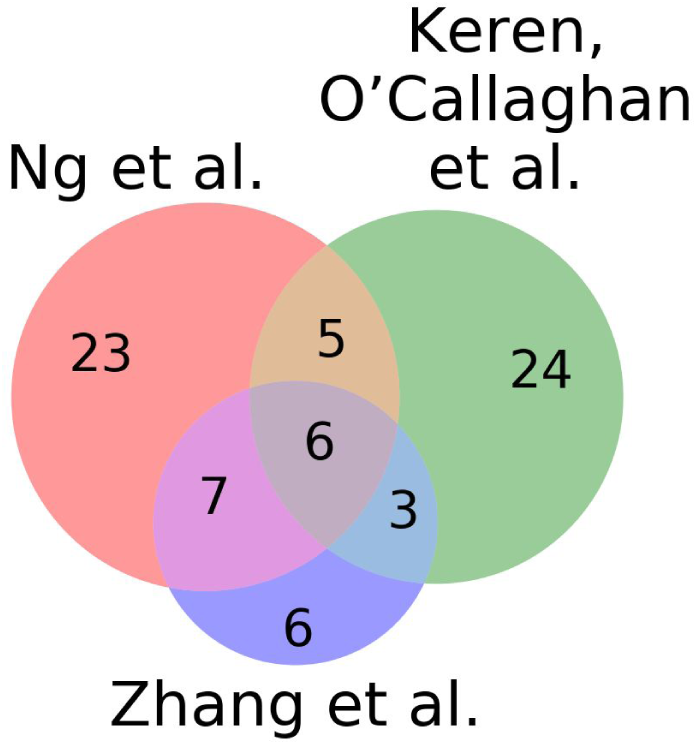
Degree of overlap in the fMRI studies reviewed by Ng *et al.* (6), Keren, O’Callaghan *et al.* (7), and Zhang *et al.* (8). The consistency of striatal findings across these studies is encouraging given the divergence in included studies.

### EEG

Keren, O’Callaghan *et al.* (7) meta-analyzed 12 studies that have compared the feedback-related negativity/reward positivity (FRN/RewP) signal between depressed and healthy participants and found a mean effect size (Cohen’s d) of 0.38 (95% CI: [0.12, 0.64]) across age ranges and a mean effect size of 0.50 (95% CI: [0.15, 0.85]) in 6 studies on children and adolescents. However, this result is unlikely to be suitable for use as a diagnostic tool. In order to give a sense of the potential discriminative capability of this association, we calculated the Area Under the Curve (AUC) for this estimate (see Salgado *et al.* (51)) and find that it corresponds to an AUC of 0.64 (95% CI: [0.54, 0.72]). An AUC of 0.64 is fairly poor for differentiating between depressed and healthy participants (0.5 is chance performance and 1 would be perfect classification). This is particularly true given that brief screening questionnaires for detection of depression, such as the 2 item Patient Health Questionnaire, have AUCs of 0.90 or 0.88 in younger subjects (52).

## Longitudinal Association

Reward processing aberrations should precede depression if they are to be a cause of it. Moreover, reward processing aberrations could be a prognostic biomarker if they predicted changes in symptoms. Here we conducted a set of pre-registered random effects meta-analyses of longitudinal studies ((19,20,23,28,35,36,53–69); https://osf.io/be4nt, https://osf.io/mp49y, https://osf.io/3dz54; see supplemental materials) to quantify the correlation between neural signals of reward processing and subsequent changes in depression symptoms (see Table S2-S5 for information extracted from these papers). We accounted for non-significant unreported effects with the MetaNSUE (70). We took the most predictive striatal or reward positivity (RewP) signal from each study. For fMRI studies we included both ROI and voxel level results with a peak in the striatum. We found that striatal fMRI signals (r: -0.10, 95% CI: [-0.17, -0.03], p: 0.0067; Fig. 4) and RewP (r: -0.17, 95% CI: [-0.29, -0.04], p = 0.0041) are both inversely related with changes in depressive symptoms in observational studies (Table 1, see Table S6 for results from treatment studies and Figures S2-S7 for additional forest plots). These estimates are upwardly biased estimates because we used the largest striatal or RewP effect from each study. We also tested a set of “global” hypotheses in which we took the strongest correlation across the entire brain from each study. We analyzed the absolute value of these correlations since we included activations, connectivity, and psychophysiological interactions. The purpose of these “global” hypotheses is to define the upper bounds of the relationship between neural reward processing signals and changes in depression symptoms. Based on this, the upper bound for the relationship is 0.17 (95% CI: [0.09, 0.24]) for observational fMRI studies, with predictions using EEG in a similar range (r: 0.20 95% CI: [0.04, 0.34]). These associations are large enough to be of mechanistic interest, but correspond to AUCs of 0.60 (95% CI: [0.55, 0.64]) for fMRI and 0.61 (95% CI: [0.50, 0.70]) for EEG and are therefore unlikely to be useful for prognosis on their own.

**Table 1:** Summary of predictive meta-analytic hypotheses. The “global” results are best-case analyses taking the absolute value of the strongest effect from any reward related analysis to define the upper bounds of the relationship between reward processing and future changes in depression. p-values are not given because significant difference from 0 is trivial after taking the absolute value. The least significant results from a leave-one-out analysis are shown in the “worst” columns.

| Modality | Specificity | Design | N | r (95% CI) | z | p | i<sup>2</sup> | Worst r | Worst z | Worst p |
| --- | --- | --- | --- | --- | --- | --- | --- | --- | --- | --- |
| fMRI | Striatum | Obs. | 9 | -0.10 [-0.17, -0.03] | -2.75 | 0.0067 | 2.80% | -0.08 | -2.26 | 0.024 |
| EEG | RewP | Obs. | 5 | -0.17 [-0.29, -0.04] | -2.63 | 0.0085 | 69.63% | -0.11 | -2.07 | 0.038 |
| fMRI | Global | Obs. | 13 | 0.17 [0.09, 0.24] | 4.30 |  | 47.86% | 0.14 | 3.76 |  |
| EEG | Global | Obs. | 5 | 0.20 [0.04, 0.34] | 2.54 |  | 79.30% | 0.12 | 2.16 |  |

**Figure 4:**
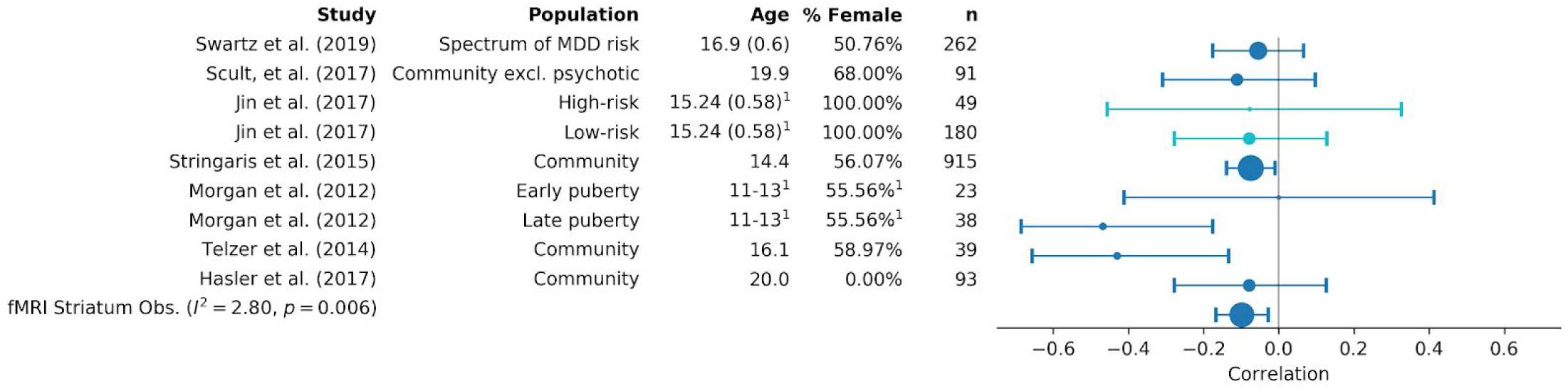
Forest plot for random effects meta-analysis of observational fmri studies reporting a striatal effect for the correlation with change in depressive symptoms. Across these studies (23,28,55,59,62,66,69), predominantly conducted in adolescents, we found that the mean of the distribution effects size for similar studies was -0.10 [-0.17, -0.03]. ^1^ indicates statistics reported for the entire study population, not for the subgroup upon which displayed prediction is based.

### Effect of measurement error on observed correlations

Our meta-analytic estimates are limited by the error in measuring neural reward signals and depressive symptoms (71), so we estimated the magnitude of the “true” relationship in the absence of measurement error and examine the implications of that estimate for future studies.

There are varied estimates for the reliability of reward processing in task-based fMRI (72–75), but overall reliability of signals from task-based fMRI is poor (42). We conducted a random effects meta-analysis across the 9 reward related fMRI analyses from Elliot *et al.* (42) (Table S7) and found the test-retest reliability to be 0.44 (95% CI: [0.28, 0.58]). Measures of depressive symptoms tend to have higher test-retest reliability. We conducted an informal review of the literature and estimated the test-retest correlation of depression measures to be 0.77 (95% CI: [0.67, 0.84]; Table S8). With some simplifying assumptions, we derived the algebraic relationship between these test-retest reliabilities (neural signals of reward processing: *r*_*NNm*_; depressive symptoms: *r*_*DDm*_), the measured relationship between neural reward signals and change in depression symptoms (*r*_*NmDm*_), and the “true” relationship between neural reward signals and change in depression symptoms (*r*_*ND*_) (derivations in supplemental methods):

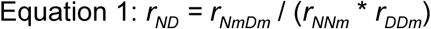

In addition, we estimated the standard error of *r*_*ND*_ with the delta method (76).

We estimated *r*_*ND*_ to be -0.29 (95% CI: [-0.57, 0.0068]), but as this is based on informal literature reviews for the test-retest reliabilities, Figure 5 shows this relationship for all values of *r*_*NNm*_ and *r*_*DDm*_ compatible with *r*_*NmDm*_ = -0.10, which is what we found for the correlation between striatal fMRI reward signal and change in depression symptoms in observational studies. A correlation of -0.29 is strong enough that striatal fMRI reward signals may be a useful predictor of future symptoms, but we would only approach that power if the reliability can be improved. If we assume that *r*_*ND*_ is -0.29, we can quantify the expected *r*_*NmDm*_ in future studies using measures with different reliabilities (Figure 6). Additionally, it is important to keep in mind that this is an optimistic estimate of the correlation between neural reward processing signals and change in depressive symptoms, and that cross-validated predictive performance would likely be lower.

**Figure 5:**
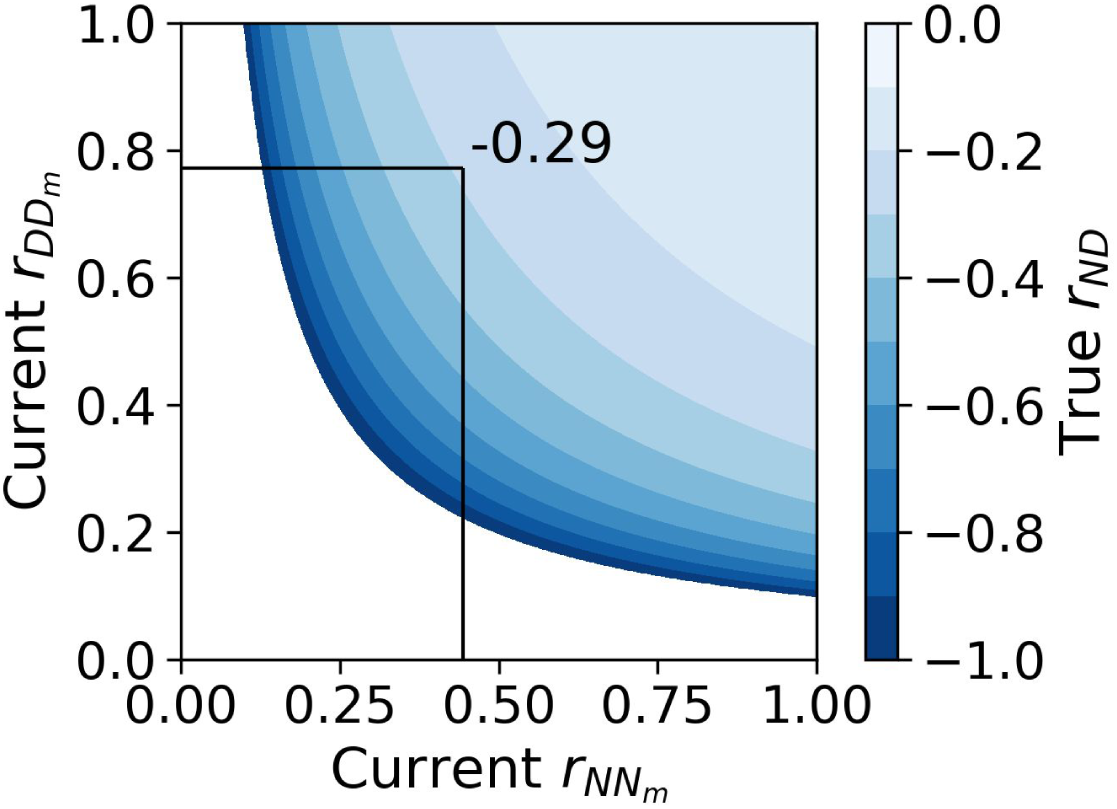
Relationship between test-retest reliabilities of depression measures (Current *r*_*DDm*_), neural reward processing signals (Current *r*_*NNm*_), and the “true” correlation between neural reward signals and change in depression symptoms (*r*_*ND*_) using our meta-analytic finding for the measured correlation between striatal fMRI reward signal and change in depression symptoms in observational studies (*r*_*NmDm*_) of -0.10. The estimate of the true effect depends on the estimates of the test-retest reliabilities, with our estimates (indicated by black lines) we find *r*_*ND*_ =-0.29 (95% CI: [-0.57, 0.0068]). Since our estimates come from informal reviews, we provide this visualization of the relationship given different values of these reliabilities. The white region indicates values of *r*_*NNm*_ and *r*_*DDm*_ incompatible with *r*_*NmDm*_ = -0.10.

**Figure 6:**
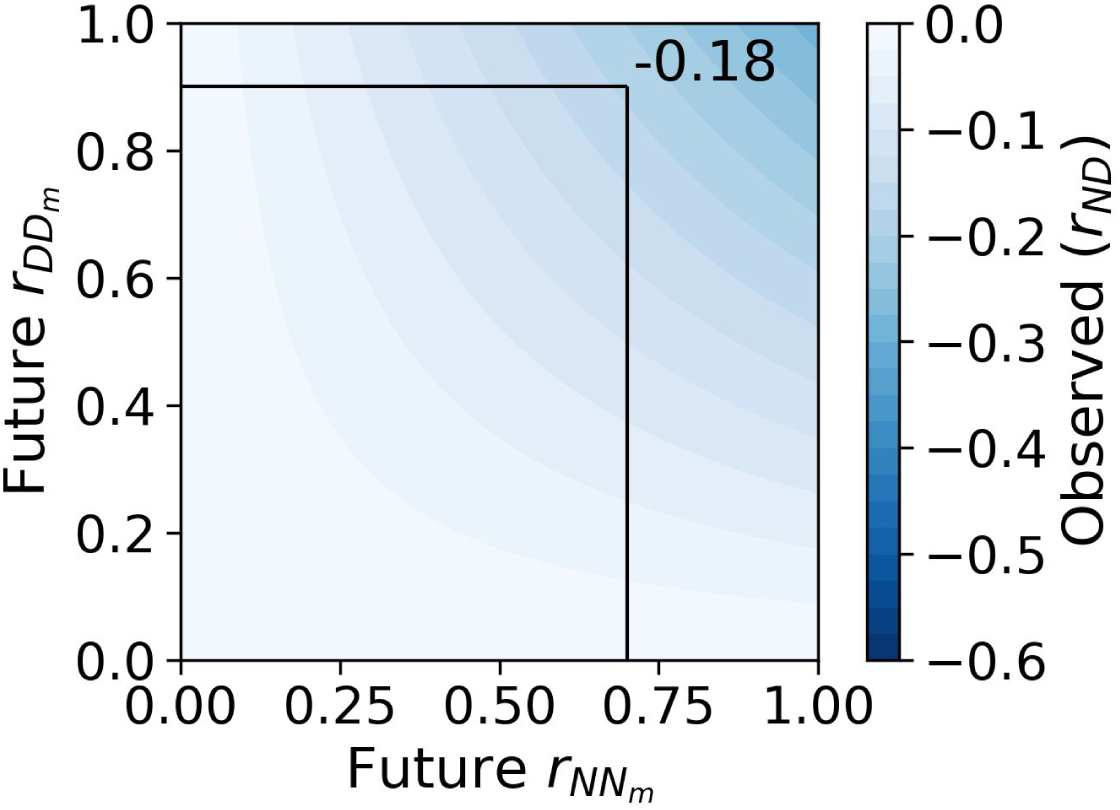
Relationship between test-retest reliabilities of depression measures (Future *r*_*DDm*_), neural reward processing signals (Future *r*_*NNm*_), and the observed correlation between neural reward signals and change in depression symptoms (*r*_*NmDm*_) using our estimate for the true correlation (*r*_*ND*_) of -0.29. We plot an illustrative example (black lines) for a hypothetical future study with *r*_*DDm*_ *= 0.9 and r*_*NNm*_ = 0.7, which gives an expected observed correlation of -0.18 (95% CI: [-0.34, 0.0043]). Researchers using our estimate in this way should keep in mind that our longitudinal meta-analysis took the best correlation between activations and depressive symptoms across the entire striatum and they should plan to account for multiple comparisons appropriately.

As pre-registered, we conducted a review of the quality of the papers included in our meta-analysis on the basis of best practices for predictions based on neuroimaging data (10) and open science practices (77). Only two studies provided cross-validated metrics for prediction accuracy (Table S1). There were four studies that referenced pre-registration (19,20,35,63), but none of these had in depth descriptions of analytical approaches.

There is evidence that reward processing signals correlate with changes in depression symptoms. While only a modest correlation is observed (r = -0.10 [-0.17, -0.03]), more reliable measures may find a larger relationship--up to unknown limits of measurement reliability. On the other hand, if taken at face value, these modest observed correlations are consistent with the hypothesized mechanism of reward processing aberrations causing anhedonia and depression. In daily life we are constantly receiving rewards, a small difference in the processing of those rewards may accumulate over time to have a significant impact on clinical symptoms (78).

## Manipulability

If reward processing abnormalities cause depression, then altering the reward processing network should alter the clinical phenotype and course of depression. Manipulating reward stimuli changes ventral striatum activity as well as subjective ratings of momentary mood (46,79). However, evidence that manipulating the reward processing system changes clinical symptoms of depression has been elusive. The ideal evidence would come from a randomized, placebo controlled trial, where the intervention would be shown to a) cause a change in reward processing; b) through that change in reward processing, cause a change in behavior. Statistically, this amounts to a mediation. Many interventions can perturb the reward system (80–89) and some have even shown that changes in this system correlate to changes in depressive symptoms (20,63,90–93). One study did find a significant mediation (20), so they provide some evidence for the manipulability of depression symptoms via manipulations of the reward processing system.

### Statistical power

Mediation designs are particularly demanding in terms of power. If we assume small effect sizes (βs around 0.1) and only two measurements then 395 subjects would be needed for 80% power if a bootstrap test of mediation is used (94). Increasing the number of measurements to five reduces the required number of subjects to 253 and may give insights into mechanisms of long delays in treatment response and differential behavioral effects during the course of treatment (95). Only one of the six intervention studies mentioned above had the 55 participants needed for 80% power to detect even moderate effect sizes, that is βs around 0.4. Please see Pan *et al.* (94) for details on determining sample size for mediation analyses.

## Conclusion

Neural reward processing abnormalities are currently unsuited for use as clinical predictors of depression, but improved measures of neural signals of reward processing and multivariate analyses may change this in the near future. There is evidence to support a causal relationship between reward processing abnormalities, with weak temporal association and evidence for manipulability. We have made general recommendations for best practices (Table 2) and specific experimental recommendations (Table 3) for addressing some of the challenges we observed in the literature. Not all of these recommendations are applicable to every study of reward processing and depression, but we hope that they will be a useful guide to the design of future studies.

**Table 2:** General recommendations for best practices.

| Challenge | Recommendation |
| --- | --- |
| *Theoretical weaknesses* | • Make strong theoretical predictions against alternative models e.g. reward anticipation vs reward learning in depression (47).<br>• Specify potential causal pathways in the relationship between parameters of reward processing and depression e.g. conduct a highly sampled longitudinal assessment of reward processing and depression. Cross-lagged analysis of the relationship between reward processing and depression will allow separation of time-varying (state) and time-invariant (trait) components.<br>• If focusing on predictive utility, use up-to-date methodology, including cross-validation (as in the responder classification analysis here (56)) or out-of-sample validation, sample sizes of at least several hundred observations, provide multiple measures of prediction accuracy, use sum-of-squares formulation for $R^2$, k-fold cross-validation with k between 5 and 10, and sharing of data sets and code (10,14,77). |
| *Developmental moderation* | • Spell out developmental theory of reward in relation to depression and build clear expectations on that basis.<br>• Use a study that spans development at least in accelerated design (96) or better in longitudinal design sensitive to developmental stages (97,98).<br>• Use well-powered study with explicit calculations for any higher order interactions, e.g. between reward processing and pubertal stage or timing.<br>• Include biological samples relevant to development, including sex hormones. |
| *Cross-sectional association* | *Employ and facilitate more powerful meta-analytic techniques:*<br>• Apply meta-analysis that are sensitive to direction and magnitude of effect (such as in Albajes-Eizagirre *et al.* (70)).<br>• Share statistical maps on platforms such as NeuroVault.org. This would allow use of more powerful image based meta-analysis techniques (99). |
| *Manipulability* | *Design studies to explore treatment and mediation effects:*<br>• Include behavioral and or clinical outcomes in treatment studies, even acute treatments (100).<br>• Make clear predictions about mediation effect (e.g. of reward processing parameter) and power study accordingly.<br>• Use multiple assessments as this allows for better estimation of timing of change and increases power. See Pan *et al.* (94) for details. |

**Table 3:** Specific experimental recommendations.

| **Challenge** | **Recommendation** |
| --- | --- |
| *Specificity* | • Compare theory of global reward processing deficit (“analogy to physical illness”) with context-specific anhedonia (27).<br>• Assess idiosyncratic effects on reward anticipation and experience and how these vary.<br>• Build and test theories about differential pathways to anhedonia, e.g. compare anhedonia in psychosis to classical depressive anhedonia (101).<br>• Use well-powered samples and designs.<br>• Consider hetero-typic continuities (e.g. social phobia to social anhedonia to depression). |
| *Difficulty measuring anhedonic symptoms and reward processing aberrations* | *Develop a reward processing task, anhedonic symptom questionnaire, and generative computational model in concert.* These should:<br>• Be developed collaboratively (similar to the model used in the development of BIDS (102)) to promote use and adoption.<br>• Behavioral tasks of anhedonia/motivation should be geared towards capturing intra-individual change with good test-retest reliability (41).<br>• Measure all or many phases of reward processing in tandem (103), ideally one that is also easy to use developmentally.<br>• Measure multiple aspects of anhedonia, ideally in a non-retrospective or vicarious way, to disentangle recall of reward from actual anticipation or experience of reward.<br>• Tasks should ideally have animal analogues to facilitate more direct translation between humans and model organisms.<br>• Explicitly link reward processing parameters with symptoms as measured by questionnaires so that theoretical links between task performance and symptoms are explicit where feasible.<br>• Be amenable to repeated administration for use in longitudinal studies. |

## Acknowledgements

All data collated for this study is available at https://osf.io/whvam/.

All code for analyses in this study are at https://github.com/nimh-mbdu/great_expectations.

This research was supported in part by the Intramural Research Program of the National Institute of Mental Health, National Institutes of Health (grant ZIA-MH002957-01 to Dr. Stringaris). The funder had no role in the design and conduct of the study; collection, management, analysis, and interpretation of the data; preparation, review, or approval of the manuscript; and decision to submit the manuscript for publication.

## Disclosures

None of the authors have any biomedical financial interests or potential conflicts of interest.

## Notes

https://osf.io/whvam/&#8203;

https://github.com/nimh-mbdu/great_expectations&#8203;

https://mybinder.org/v2/gh/nimh-mbdu/great_expectations/de3c6434b364f76db5b7e4b31f218d40459e3c0e

