## Supplemental Materials for "Great Expectations: A Critical Review of and Recommendations for the study of Reward Processing as a Cause and Predictor of Depression"

**Supplemental Methods**

Preregistrations and deviations

We pregisterred the approach to our literature review (https://osf.io/be4nt, https://osf.io/mp49y) and meta-analysis (https://osf.io/3dz54). The first registration (https://osf.io/be4nt) described our methodology, repositories, search terms, and inclusion/exclusion criteria. The second was a note modifying the search period from going through the end of 2019 to stopping at December, 17th, 2019 due to the limited search granularity provided by many of the publication databases we used. We added a second round of title and abstract screening in addition to the single round specified in these registrations and had two separate investigators (AS and DMN) review articles for inclusion or exclusion.
The last registration (https://osf.io/3dz54) is for the analytical and qualitative approach for the meta-analyses. This registration was completed after reviewing papers for inclusion, but prior to beginning data extraction for the meta-analysis. In our registration we referred to feedback negativity, which is the subtraction of gain ERP from loss ERP. In reviewing the studies, we found that reward positivity (RewP, the subtraction of loss ERP from gain ERP) was more commonly reported and so we use RewP here. In our registration we stated that we would only use MetaNSUE if non-significant unreported effects made up 20% or more of the relevant effects to a particular hypothesis, but we instead used it for all analyses for consistency.

Meta-analysis

*Literature review:* We searched PubMed, Scopus, PsycINFO, and Web of Science for articles published in English from February 1, 2017 to December 17, 2019, using the following terms and their derivatives: depression, anhedonia, reward, motivation, reinforcement, punishment and aversion, prediction error, decision making, and risk taking. This is the specific query:
(((((depress*) OR (anhedon*)) AND ((reward*) OR (motiv*) OR (reinforc*) OR (punish*) OR (aversi*) OR ("prediction error") OR ("decision making") OR ("risk taking")))) AND ("2017/02/01"[Date - Publication]: "2019/12/31"[Date - Publication])) AND English[Language]
Studies returned by this initial query were reduced by removing duplicates and non-human and non-experimental studies based on keyword searches. 30 articles were randomly selected from the remainder and screened for inclusion/exclusion by independent investigators (DMN, CCC, SK, CW, SMJ, LG). The inter-rater reliability was found to be 0.88 and the remaining sample of studies was divided among the investigators for abstract screening. Each article was screened twice and those that passed either screening were divided up among the investigators for a more thorough review by two investigators (AS and DMN). These two investigators decided together which articles to include and exclude.

*Inclusion criteria:* To be included, studies had to provide a measure of depression or anhedonia in people with major depressive disorder, in people at high risk of depression, or in healthy volunteers. We selected only studies that measured depression or depressive symptoms through questionnaires, structured interviews, or clinical diagnosis. In terms of reward paradigms employed, and following the classification described by Richards et al. (1), we included instrumental- reward tasks and decision-making tasks, which require participants to complete an action correctly in order to obtain a reward, as this action is linked to the reward value at a trial- by-trial level. Hence, reward paradigms in which rewards were presented passively were excluded. Either positive (e.g., winning money) or negative (e.g., losing money) reward manipulations were permitted. No age restrictions were applied. In addition, studies must have reported analyses examining the ability of measures collected at one time point to predict severity (as evidenced by changes in depression/anhedonia status or scores) or treatment response at a subsequent time point.

*Exclusion criteria:* Studies were excluded if they lacked a standard measure of depression. We excluded studies that measured depressive symptoms in patients with another disorder (e.g. bipolar or schizophrenia, etc.), but did not include in addition a depressed group. This was done because our primary question concerns the effects of depression on reward processing and in the absence of a depressed control group, drawing inferences about such effects would be impossible. We also excluded studies in which reward processing was only measured through non-experimental methods such as self-report measures or questionnaires. Furthermore, to guard against heterogeneity, we excluded studies in which physical punishment was delivered (e.g. heat, pain, electrical shock, etc.) as these are likely to engage different brain networks. Similarly, studies were excluded if they employed passive exposure to pleasant/unpleasant stimuli such as facial emotions or images. For included studies, relevant methodological details, where available, were recorded, as outlined below.

*Data Extraction:* We only evaluated one prediction for each set of study participants. If multiple predictions were reported for the same set of subjects we picked the prediction over the longest time period, if multiple predictions were made with the same time period, we picked the highest quality prediction which used the most participants. Each study could potentially provide more than one prediction if they reported separately on different samples.
We rated the quality of each longitudinal prediction based on the criteria put forward in Poldrack et al. (2). Specifically:

- Reporting of out of sample model fit indices
  - Least risk for a completely separate evaluation set. Highest risk for in-sample model fit indices reported as predictive performance.
    - 0: completely separate test set
    - 1: within sample cross validation
    - 2: in-sample model fit reported as predictive performance
- Cross-validation procedure that encompasses all analytical manipulations
  - Highest marks if all normalizing and selection done separately for each validation fold. Lowest if only the final model fit is done in a cross validated fashion.
    - 0: all cross-dataset analytical procedures performed within training set
    - 1: some analytical procedure performed within the training set, but not all
    - 2: significant chance of data leakage between training and test
- Sample size
  - Ideally greater than several hundred observations
    - 0: > 200
    - 1: > 50 < 200
    - 2: <50
- Reporting multiple measures of model fit
  - Lowest risk if multiple measures such as $R^2$ in addition to measures of unsigned error like mean square error or mean absolute error are provided. Highest risk if none of these measures is provided.
    - 0: provided at least 2 measures of predictive performance, one of which is MSE or absolute error
    - 1: only r-squared or a single metric reported

        - 2: no standardized predictive accuracy reported
- Calculating coefficient of determination via sum-of-squares formulation
    - Lowest risk for sum-of-squares formulation. Highest risk for squared correlation coefficient.
        - 0: sum-of-squares formulation
        - 2: squared correlation coefficient
- Use of k-fold or shuffle-split cross validation
    - Lowest risk for shuffle-split or k-fold cross validation with k between 5 and 10. Highest risk for leave-one-out cross-validation.
        - 0: K-fold or shuffle split cross validation with k between 5 and 10
        - 1: some intermediate cross validation scheme
        - 2: leave one out cross validation

In addition, we evaluated the open science practices of the publication on the following criteria:

- Use of pre-registration
    - 0: if a link to a thorough pre-registration is provided
    - 1: if link is provided but there are extensive deviations from the prregistration
    - 2: no pre-registration
- Publicly releasing code or scripts used in analysis
    - 0: link to working code provided
    - 1: link provided, but code does not work
    - 2: no link to code
- Publicly releasing data used in analysis
    - 0: raw data available with or without a data-use-agreement
    - 1: statistical maps or other derivatives released
    - 2: no data released

Each criteria was rated on a scale of 0 for least risk of bias to 2 for high risk of bias as described above, or N/A, for example the K-fold criteria is not applicable if no cross-validation was performed. Quality was evaluated by two separate raters (DMN and GO'C), conflicts were reconciled in person and are reported in.

Two raters (DMN and GO'C) independently extracted the information described below from each longitudinal prediction.

- Observational or treatment
- Study in which source data were first described
- Nature of each group (healthy, at risk (defined as the presence of either MDD in a parent, high depression scale scores in the absence of MDD diagnosis, or remitted MDD, depressed), participants with another disorder)
- Criteria used for diagnosis if relevant
- Sample size of each group
- Percentage of females in each group
- Percentage of medicated individuals in each group
- Mean, SD, and range of ages in each group
- Neural measure: EEG, functional connectivity, fMRI
- Depression measure
- Reward task
- Type of reward (monetary, affective, or primary)
- Contrast used (if any)
- Prediction interval in time

- Terms used in the predictive model
- Link to preregistration
- Link to code
- Link to data
- Treatment type (pharmacological or psychological)
- Specific treatment

For fMRI studies we separately extracted information for the location/connection providing the best predictive performance across the entire brain and the striatal location/connection with the best predictive performance:

- Direction of effect
- Reported statistic
- Predictive effect size uniquely contributed by the neural information
- Overall predictive effect size
- Template space of reported components
- Coordinates of effect
- Connections

For EEG studies extracted information for both the most predictive component of the EEG signal and for the RewP:

- Electrodes sampled from
- Type of signal extracted (FRN, RewP, etc.)
- Sampling method (mean amplitude, peak, etc.)
- Window for sampling
- High and low pass filter applied
- Reference electrode
- Type of EEG net used
- Direction of effect
- Reported statistic
- Predictive effect size uniquely contributed by the neural information
- Overall predictive effect size

Reported effects were transformed to have a uniform direction of effect as appropriate, for example, the sign of feedback negativity results were flipped so that they were in terms of RewP. Reported effects were transformed into correlation coefficients reflecting the unique contribution of neural data. Transformations from t or F statistics to correlation coefficients based on the following formulas (3):

$$r = \sqrt{\frac{t^2}{(t^2 + df)}}$$

$$r = \sqrt{\frac{df_n F}{df_n F + df_d}}$$

*df*: degrees of freedom; $df_n$: numerator degrees of freedom; $df_d$: denominator degrees of freedom
When only a beta coefficient was reported, this was converted to a t by dividing by the standard error and then from t to correlation coefficient.

*Analytic approach:* We sought to test the central hypothesis that the expected effect size for a study predicting future depression severity from neural reward related signals is not 0. We operationalized this hypothesis into 8 separate hypotheses based on the modality (fMRI or EEG), specificity (striatal reward contrast/RewP or any signal), and study design (treatment or observational). We established an a priori minimum sample size of 5 predictions for each of these hypotheses. We performed random-effects meta-analysis of correlation coefficients using Fisher's transform with multiple imputation of non-significant unreported effects as implemented in the R package MetaNSUE (4,5). We rely on the tables converting effects sizes to area under the receiver operator characteristics curve (AUC) provided by Salgado (6) to help put the effect sizes we report in a form more familiar to machine learning practitioners. Correlation coefficients are first converted to d before converting to AUC.

Accounting for the effect of measurement error on observed correlations

After completing our meta-analysis, we wanted to understand how our findings constrained the distribution of true values for the correlation between neural reward processing signals and future depressive symptoms given that both depressive symptoms and reward processing are subject to measurement error. We first estimated the test-retest reliability of neural reward signals and depressive symptoms, then we used the algebraic relationships derived below to calculate the true relationship between neural reward processing signals and future depressive symptoms based on the meta-analytic estimate of correlation between striatal fMRI reward signal and change in depression symptoms in observational studies. Finally, we reversed the calculation to determine how measurement error in future studies would impact the observed effect size.

*Estimating measurement error of neural signals:* We extracted the intra class correlations, sample sizes, and measurement intervals from Elliott et al. (7) (Supplemental Table 7) for reward related tasks with a prediction interval of less than 100 days. This resulted in 9 values from 7 studies from which to meta-analytically estimate the reliability of neuroimaging measures of reward. We then ran a random effects meta-analysis to determine the mean correlation, standard error of that estimate, and an estimate of the between studies heterogeneity, $\tau^2$.

*Estimating measurement error of depression symptoms:* We conducted an informal literature review to identify studies that assessed test-retest reliability of the measures of depression used in the studies in our longitudinal meta-analysis. We extracted the correlation coefficient, sample size, prediction interval, and description of the population from studies that examined test-retest reliability of depression measures over an interval of less than 100 days (Supplemental Table 8). This gave us 13 reliability estimates for 6 measures from 6 studies. We then ran a random effects meta-analysis to determine the mean correlation, standard error of that estimate, and an estimate of the between studies heterogeneity, $\tau^2$.

*Deriving relationship between measured and true correlation:* We algebraically derived the relationship between between the above test-retest reliabilities (neural signals of reward processing: $r_{NNm}$; depressive symptoms: $r_{DDm}$), the measured relationship between neural reward signals and change in depression symptoms ($r_{NmDm}$), and the "true" relationship between neural reward signals and change in depression symptoms ($r_{ND}$). In order to do so, we assume an additive model for noise, such that the covariance between the underlying constructs and their measurement is 1 and the standard deviation of the underlying construct is 1. This simplifies the relationship between test retest correlation and the standard deviation of the measured properties. Formally:

$$r_{NN_m} = \frac{cov(N, N_m)}{\sigma_N \sigma_{N_m}}$$

Assume $\sigma_N = 1$, $cov(N, N_m) = 1$

$$r_{NN_m} = \frac{1}{\sigma_{N_m}}$$

$$r_{DD_m} = \frac{cov(D, D_m)}{\sigma_D \sigma_{D_m}}$$

Assume $\sigma_D = 1$, $cov(D, D_m) = 1$

$$r_{DD_m} = \frac{1}{\sigma_{D_m}}$$

Given that we've assumed an additive noise model, we assume that the covariance between measured depressive symptoms and measured neural measures of reward processing is the same as the covariance between the true values. With these simplifications in place the relationship between measured correlation and observed correlation is greatly simplified.

$$r_{ND} = \frac{cov(ND)}{\sigma_N \sigma_D}$$

Since $\sigma_N = 1$ and $\sigma_D = 1$

$$r_{ND} = cov(ND)$$

$$r_{N_m D_m} = \frac{cov(N_m D_m)}{\sigma_{N_m} \sigma D_m}$$

Assume $cov(N_m D_m) = cov(ND) = r_{ND}$

$$r_{N_m D_m} = r_{ND} \cdot r_{NN_m} \cdot r_{DD_m}$$

$$r_{ND} = \frac{r_{N_m D_m}}{r_{NN_m} \cdot r_{DD_m}}$$

While these assumptions are necessary to make this estimation tractable, they do represent an optimistic scenario.

With these equations in hand, we can use the delta rule (8) to combine the estimated standard errors across the three measured correlations and derive confidence intervals for our estimate of $r_{ND}$. Specifically:

$$\sigma_{r_{ND}} = \sqrt{r_{ND} \left(\frac{\sigma_{r_{NN_m}}}{r_{NN_m}}\right)^2 \left(\frac{\sigma_{r_{DD_m}}}{r_{DD_m}}\right)^2}$$

And the confidence interval is derived from the standard error with a Bonferroni correction for the number of estimated values, giving a critical z-score of 2.39. When estimating $r_{N_m D_m}$ for some future study, we have point estimates for the test-retest reliabilities so we multiply the limits of the confidence interval for $r_{ND}$ by the test-retest-reliabilities to obtain the confidence interval of future $r_{N_m D_m}$.

We calculated this relationship between the test-retest reliabilities and $r_{ND}$ across a grid of 1000 values between 0 and 1 for $r_{NN_m}$ and 1000 values between 0 and 1 for $r_{DD_m}$ to estimate $r_{ND}$ with different assumptions about the current test retest reliability of measures of depression and neural reward signals (Figure 5 of the main text). We took the values for those reliabilities that we estimated as described above and our meta-analytic estimate of correlation between striatal fMRI reward signal and change in depression symptoms in observational studies to determine our best estimate of $r_{ND}$. Assuming that best estimate is correct, we then calculated what the observed $r_{N_m D_m}$ would be for some future study across a grid of a grid of 1000 values between 0 and 1 for $r_{NN_m}$ and 1000 values between 0 and 1 for $r_{DD_m}$ (Figure 6 of the main text).

### Supplemental Tables

| Study | Zhang et al. | Keren, O'Callaghan et al. | Ng et al. |
|---|---|---|---|
| Admon et al. (9) | 0 | 1 | 0 |
| Arrondo et al. (10) | 0 | 1 | 1 |
| Bremner et al. (11) | 0 | 0 | 1 |
| Burger et al. (12) | 0 | 0 | 1 |
| Canli et al. (13) | 1 | 0 | 0 |
| Casement et al. (14) | 0 | 1 | 0 |
| Chan et al. (15) | 0 | 1 | 0 |
| Chandrasekhar Pammi et al. (16) | 0 | 1 | 0 |
| Chantiluke et al. (17) | 1 | 0 | 1 |
| Chase et al. (18) | 0 | 0 | 1 |
| Chung & Barch (19) | 0 | 1 | 0 |
| Demenescu et al. (20) | 0 | 0 | 1 |
| Derntl et al. (21) | 1 | 0 | 0 |
| Dichter et al. (22) | 1 | 1 | 1 |
| Dillon et al. (23) | 0 | 1 | 0 |
| Elliott et al. (24) | 0 | 0 | 1 |
| Engelmann et al. (25) | 0 | 0 | 1 |
| Epstein et al. (26) | 1 | 0 | 0 |
| Felder et al. (27) | 0 | 1 | 0 |
| Forbes et al. (28) | 1 | 1 | 0 |
| Forbes et al. (29) | 0 | 1 | 0 |
| Fournier et al. (2013) | 1 | 0 | 1 |
| Fu et al. (30) | 0 | 0 | 1 |
| Fu et al. (31) | 1 | 0 | 1 |
| Fu et al. (32) | 0 | 0 | 1 |
| Gorka et al. (33) | 0 | 1 | 0 |
| Gotlib et al. (34) | 1 | 0 | 1 |
| Gotlib et al. (35) | 0 | 1 | 0 |
| Gradin et al. (36) | 1 | 1 | 0 |
| Gradin et al. (37) | 0 | 0 | 1 |
| Hagele et al. (38) | 0 | 1 | 0 |

| | | | |
|---|---|---|---|
| Hall et al. (39) | 0 | 0 | 1 |
| Johnston et al. (40) | 0 | 1 | 1 |
| Keedwell et al. (41) | 1 | 0 | 1 |
| Knutson et al. (42) | 1 | 1 | 1 |
| Kumar et al. (43) | 1 | 0 | 0 |
| Kumari et al. (44) | 1 | 0 | 1 |
| Laurent et al. (45) | 0 | 0 | 1 |
| Liu et al. (46) | 0 | 0 | 1 |
| Luking et al. (47) | 0 | 1 | 0 |
| McCabe et al. (48) | 1 | 0 | 0 |
| Mitterschiffthaler et al. (49) | 1 | 0 | 0 |
| Mori et al. (50) | 0 | 1 | 0 |
| Murrough et al. (51) | 0 | 0 | 1 |
| Olino et al. (52) | 0 | 1 | 0 |
| Olino et al. (53) | 0 | 1 | 0 |
| Pizzagalli et al. (54) | 1 | 1 | 1 |
| Redlich et al. (55) | 0 | 1 | 0 |
| Remijnse et al. (56) | 1 | 1 | 1 |
| Rizvi et al. (57) | 0 | 0 | 1 |
| Robinson et al. (58) | 1 | 1 | 0 |
| Rosenblau et al. (59) | 0 | 0 | 1 |
| Rzepa et al. (60) | 0 | 1 | 0 |
| Satterthwaite et al. (61) | 0 | 1 | 0 |
| Scheuerecker et al. (62) | 0 | 0 | 1 |
| Schiller et al. (63) | 0 | 1 | 1 |
| Segarra et al. (64) | 0 | 1 | 1 |
| Sharp et al. (65) | 0 | 1 | 1 |
| Smoski et al. (66) | 1 | 1 | 1 |
| Smoski et al. (67) | 1 | 1 | 1 |
| Steele et al. (68) | 0 | 1 | 0 |
| Stoy et al. (69) | 0 | 1 | 0 |
| Stringaris et al. (70) | 0 | 1 | 0 |
| Surguladze et al. (71) | 1 | 0 | 1 |
| Surguladze et al. (72) | 0 | 0 | 1 |
| Townsend et al. (73) | 0 | 0 | 1 |
| Ubl et al. (74) | 0 | 1 | 0 |
| Ubl et al. (75) | 0 | 1 | 0 |
| Wagner et al. (76) | 0 | 0 | 1 |

| | | | |
|---|---|---|---|
| Wang et al. (77) | 0 | 0 | 1 |
| Yang et al. (78) | 0 | 1 | 0 |
| Young et al. (79) | 0 | 0 | 1 |
| Zhang et al. (80) | 0 | 0 | 1 |
| Zhong et al. (81) | 0 | 0 | 1 |

Table S1: Set of reward processing studies included across Zhang et al. (82), Keren, O'Callaghan et al. (83), and Ng et al. (84). 1 indicates a study was included in that review, 0 indicates it was not included.

| **Study** | **Hypothesis** | **Population** | **Age** | **Female** | **N** | **Mean Interval** | **Treatment** |
|---|---|---|---|---|---|---|---|
| Admon et al. (9) | fMRI Global Treat | MDD | Adult | 50.00%[1] | 14 | 84 | SAMe (5), escitalopram (5), placebo (4) |
| Bakker et al. (85) | fMRI Global Treat | Low-moderate risk | 20.9 (2.1) | 82.76% | 87 | 7.5 | reward anticipation on activity pleasantness |
| Bakker et al. (85) | fMRI Global Obs. | Low-moderate risk | 20.9 (2.1) | 82.76% | 87 | 7.5 | reward anticipation on activity pleasantness |
| Barch et al. (86) | EEG Global Treat | MDD | 5.5 (0.8) | 35.00% | 60 | 126 | Parent-Child Interaction Therapy |
| Barch et al. (86) | EEG RewP Treat | MDD | 5.5 (0.8) | 35.00% | 60 | 126 | Parent-Child Interaction Therapy |
| Bertocci et al. (87) | fMRI Global Obs. | Parent with BD or Axis-1 | 14.0 (2.3) | 46.34% | 41 | 29.6 | |
| Bress et al. (88) | EEG RewP Obs., EEG Global Obs. | HV enriched for parental MDD | 16 | 100.00% | 61 | 630 | |
| Burani et al. (89) | EEG RewP Treat, EEG Global Treat | Community | 12.6 (1.7) | 100.00% | 183 | 365.25 | sleep and stress |
| Burani et al. (89) | EEG RewP Obs., EEG Global Obs. | Community | 12.6 (1.7) | 100.00% | 183 | 365.25 | sleep and stress |
| Burkhouse et al. (90) CBT | EEG RewP Treat, EEG Global Treat | R-DOC internalizing symptoms | 28.7 (8.9) | 73.50% | 34 | 84 | CBT |
| Burkhouse et al. (90) Sertraline | EEG RewP Treat, EEG Global Treat | R-DOC internalizing symptoms | 24.9 (8.1) | 75.90% | 29 | 84 | sertraline |
| Flores et al. (91) | fMRI Global Obs. | HV | 16.3 (1.5) | 65.00% | 34 | 7 | |
| Greenberg et al. (92) | fMRI Global Treat | MDD | 36.9 (12.8) | 66.95% | 194 | 56 | sertraline or placebo |
| Greenberg et al. (92) | fMRI Global Obs. | MDD | 36.9 (12.8) | 66.95% | 200 | 56 | sertraline or placebo |
| Hasler et al. (93) | fMRI Striatum Obs., fMRI Global Obs. | Community | 20 | 0.00% | 93 | 730.5 | |

| | | | | | | | |
|---|---|---|---|---|---|---|---|
| Jin et al. (94) High-risk | fMRI Global Obs. | High-risk | 15.24 (0.58)[1] | 100.00% | 49 | 270 | |
| Jin et al. (94) High-risk | fMRI Striatum Obs. | High-risk | 15.24 (0.58)[1] | 100.00% | 49 | 270 | |
| Jin et al. (94) Low-risk | fMRI Global Obs. | Low-risk | 15.24 (0.58)[1] | 100.00% | 180 | 270 | |
| Jin et al. (94) Low-risk | fMRI Striatum Obs. | Low-risk | 15.24 (0.58)[1] | 100.00% | 180 | 270 | |
| Kujawa et al. (95) | EEG RewP Treat, EEG Global Treat | Community | 9 | 43.90% | 369 | 1095.75 | maternal depression |
| Kujawa et al. (95) | EEG RewP Obs., EEG Global Obs. | Community | 9 | 43.90% | 369 | 1095.75 | maternal depression |
| Kujawa et al. (96) | EEG RewP Treat, EEG Global Treat | Anxiety disorder + comorbidity | 13.1 (4.0) | 40.70% | 22 | 84 | CBT or sertraline |
| Langenecker et al. (97) | fMRI Global Treat | MDD | 28.1 (9.9) | 50.00% | 10 | 84 | duloxetine |
| Langenecker et al. (97) | fMRI Striatum Treat | MDD | 28.1 (9.9) | 50.00% | 10 | 84 | duloxetine |
| Luo et al. (98) | EEG RewP Obs. | HV | 18.9 (0.2) | 48.00% | 25 | 180 | |
| Luo et al. (98) | EEG Global Obs. | HV | 18.9 (0.2) | 48.00% | 23 | 180 | |
| Mackin et al. (99) | EEG RewP Obs., EEG Global Obs. | HV | 14.4 (0.6) | 100.00% | 467 | 540 | |
| Morgan et al. (100) Early Puberty | fMRI Striatum Obs., fMRI Global Obs. | Early puberty | 11-13[1] | 55.56%[1] | 23 | 730.5 | |
| Morgan et al. (100) Late Puberty | fMRI Striatum Obs., fMRI Global Obs. | Late puberty | 11-13[1] | 55.56%[1] | 38 | 730.5 | |
| Queirazza et al. (101) | fMRI Global Treat | MDD | 39.2 (12.9) | 48.65% | 26 | 90 | computerized CBT |
| Queirazza et al. (101) | fMRI Striatum Treat | MDD | 39.2 (12.9) | 48.65% | 26 | 90 | computerized CBT |
| Scult, et al. (102) | fMRI Striatum Obs., fMRI Global Obs. | Community excl. psychotic | 19.9 | 68.00% | 91 | 210 | |
| Stringaris et al. (70) | fMRI Striatum Obs., fMRI Global Obs. | Community | 14.4 | 56.07% | 915 | 730.5 | |
| Swartz et al. (103) | fMRI Global Obs. | Spectrum of MDD risk | 16.9 (0.6) | 50.76% | 262 | 365.25 | |
| Swartz et al. (103) | fMRI Striatum Obs. | Spectrum of MDD risk | 16.9 (0.6) | 50.76% | 262 | 365.25 | |
| Telzer et al. (104) | fMRI Striatum Obs., fMRI Global Obs. | Community | 16.1 | 58.97% | 39 | 365.25 | |
| Walsh et al. (105) | fMRI Global Treat | MDD | 33.0 (7.1) | 71.05% | 186 | 52.50[2] | BA |

Table S2: Demographic information from longitudinal studies. BA: Behavioral Analysis; CBT: Cognitive Behavioral Therapy; MDD: Major Depressive Disorder

[1]indicates statistics reported for the entire study population, not for the subgroup upon which displayed prediction is based.

[2]slope of biweekly assessments over course of 15 weeks assessed with mixed effects model

| **Study** | **Reward Type** | **Task** | **Contrast** | **ROI** | **Statistic** | **Value** | **r** |
|---|---|---|---|---|---|---|---|

| | | | | | | | |
|---|---|---|---|---|---|---|---|
| Admon et al. (9) | monetary | MID | dACC-caudate connectivity during gain - dACC-caudate connectivity during loses | caudate | r | 0.56 | 0.56 |
| Bakker et al. (85) | monetary | reinforcement learning | reward prediction error | right putamen | b | 0.05 | 0.32 |
| Bakker et al. (85) | monetary | reinforcement learning | reward prediction error | right putamen | b | 0.19 | 0.19 |
| Barch et al. (86) | points | doors | RewP | | t | 2.09 | 0.26 |
| Barch et al. (86) | points | doors | RewP | | NSUE | | |
| Bertocci et al. (87) | monetary | card guessing task | gain - neutral | mean beta from 15 significant clusters across the brain, no striatal clusters | NSUE | | |
| Bress et al. (88) | monetary | doors with concurrent negative mood induction | FN | | F | -5.46 | -0.46 |
| Burani et al. (89) | monetary | doors | RewP | | B | 0 | -0.22 |
| Burani et al. (89) | monetary | doors | RewP | | B | -0.2 | -0.06 |
| Burkhouse et al. (90) CBT | monetary | guessing | RewP | | t | -2.04 | -0.33 |
| Burkhouse et al. (90) Setraline | monetary | guessing | RewP | | t | 0.75 | 0.14 |
| Flores et al. (91) | social | social reward | high positive - neutral | right posterior superior temporal sulcus/ temporoparietal junction | r | 0.48 | 0.48 |
| Greenberg et al. (92) | monetary | card guessing task | reward index | left ventral striatum | F | 12.93 | 0.25 |
| Greenberg et al. (92) | monetary | card guessing task | reward index | right orbitofrontal cortex | F | 6.28 | 0.17 |
| Hasler et al. (93) | monetary | card guessing task | gain - baseline | ventral striatum | r | -0.08 | -0.08 |
| Jin et al. (94) High-risk | monetary | doors | loss - baseline | OFC | r | -0.37 | -0.37 |
| Jin et al. (94) High-risk | monetary | doors | loss - baseline | striatum | NSUE | | |
| Jin et al. (94) Low-risk | monetary | doors | loss - baseline | OFC | r | 0.02 | 0.02 |
| Jin et al. (94) Low-risk | monetary | doors | loss - baseline | striatum | NSUE | | |
| Kujawa et al. (95) | monetary | doors | RewP | | b | -0.12 | -0.1 |
| Kujawa et al. (95) | monetary | doors | RewP | | b | -0.07 | -0.12 |
| Kujawa et al. (96) | monetary | doors | RewP gains | | t | -2.1 | -0.42 |
| Langenecker et al. (97) | monetary | MID | gain - neutral | right inferior frontal gyrus | z | -3.53 | |

| | | | | | | | |
|---|---|---|---|---|---|---|---|
| Langenecker et al. (97) | monetary | MID | gain - neutral | putamen | z | -3.1 | |
| Luo et al. (98) | monetary | MID with self and charitable outcomes | FRN | | t | -1.27 | -0.25 |
| Luo et al. (98) | monetary | MID with self and charitable outcomes | eudaimonic anticipation vs neutral | | F | 5.36 | 0.44 |
| Mackin et al. (99) | monetary | doors | RewP | | b | -0.1 | -0.12 |
| Morgan et al. (100) Early Puberty | monetary | card guessing task | reward anticipation - baseline | | r | 0 | 0 |
| Morgan et al. (100) Late Puberty | monetary | card guessing task | reward anticipation - baseline | caudate | t | -3.23 | -0.47 |
| Queirazza et al. (101) | points | Probabilistic reversal-learning task | parametric weighed RPE | cluster in right amygdala and right hippocampus | r | -0.64 | -0.64 |
| Queirazza et al. (101) | points | Probabilistic reversal-learning task | parametric weighed RPE | cluster in right putamen and caudate | r | -0.56 | -0.56 |
| Scult, et al. (102) | monetary | card guessing task | positive feedback > negative feedback | bilateral ventral striatum | b | -0.09 | -0.11 |
| Stringaris et al. (70) | monetary | MID | anticipation of large win versus anticipation of no win | left ventral striatum | t | -2.28 | -0.08 |
| Swartz et al. (103) | monetary | MID | gain - neutral anticipation | mean of bilateral ventral striatum small volume corrected clusters with significant activation | B | 4.17 | 0.16 |
| Swartz et al. (103) | monetary | MID | gain - neutral anticipation | mean of bilateral ventral striatum small volume corrected clusters with significant activation | B | -0.83 | -0.06 |
| Telzer et al. (104) | monetary | family donation task | Costly donation > control | ventral striatum | B | -5.3 | -0.43 |
| Walsh et al. (105) | monetary | MID | gain - neutral | right putamen | t | 2.82 | 0.2 |

Table S3: Prediction information for longitudinal studies. OFC: Orbitofrontal Cortex; FRN: Feedback Related Negativity; RewP: Reward Positivity; dACC: dorsal Anterior Cingulate Cortex; MID: Monetary Incentive Delay task; NSUE: Non-Significant Unreported Effect

| Study | Out of Sample | Comprehensive CV | Sample Size | N | Multiple fit metrics | Fit Metrics | $R^2$ method | CV Method |
|---|---|---|---|---|---|---|---|---|
| Scult, et al. (102) | 2 | NA | 1 | 91 | 2 | B only | NA | NA |
| Jin et al. (94) High-risk | 1[1] | 0 | 2 | 49 | 1 | r | 2 | 0 |
| Jin et al. (94) Low-risk | 1[1] | 0 | 1 | 180 | 1 | r | 2 | 0 |
| Flores et al. (91) | 2 | NA | 2 | 34 | 1 | r | 2 | NA |
| Burkhouse et al. (90) CBT | 2 | NA | 2 | 34 | 2 | B, t | NA | NA |
| Burkhouse et al. (90) Setraline | 2 | NA | 2 | 29 | 2 | B, t | NA | NA |
| Bakker et al. (85) | 2 | NA | 1 | 87 | 2 | β | NA | NA |
| Luo et al. (98) | 2 | NA | 2 | 25 | 1 | B, t, $R^2$ | 2 | NA |
| Luo et al. (98) | 2 | NA | 2 | 25 | 2 | B, t | NA | NA |
| Barch et al. (86) | 2 | NA | 1 | 60 | 2 | B, t | NA | NA |
| Barch et al. (86) | 2 | NA | 2 | 44 | 1 | B, t | NA | NA |
| Kujawa et al. (95) | 2 | NA | 0 | 369 | 2 | β | NA | NA |
| Langenecker et al. (97) | 2 | NA | 2 | 10 | 2 | Voxel Z | NA | NA |
| Bertocci et al. (87) | 1[2] | 1[5] | 1 | 55 | 0 | β, Sum of squared error | NA | 1[6] |
| Swartz et al. (103) | 2 | NA | 0 | 262 | 2 | B | NA | NA |
| Goldstein et al. (106) | 2 | NA | 0 | 369 | 2 | B, t | NA | NA |
| Burani et al. (89) | 2[3] | NA | 1 | 183 | 2 | B | NA | NA |
| Mackin et al. (99) | 2 | NA | 0 | 467 | 2 | B | NA | NA |
| Kujawa et al. (96) | 2 | NA | 2 | 27 | 2 | B, t | NA | NA |
| Walsh et al. (105) | 2 | NA | 2 | 38 | 1 | t, pseudo $R^2$ | 2 | NA |
| Queirazza et al. (101) | 2[4] | NA | 2 | 37 | 1 | r | 2 | NA |
| Greenberg et al. (92) | 2 | NA | 0 | 222 | 2 | F | NA | NA |
| Admon et al. (9) | 2 | NA | 2 | 14 | 1 | ΔF, $\Delta R^2$ | 2 | NA |
| Stringaris et al. (70) | 2 | NA | 0 | 915 | 2 | β | NA | NA |
| Bress et al. (88) | 2 | NA | 1 | 68 | 2 | F, β | NA | NA |
| Morgan et al. (100) Early Puberty | 2 | NA | 2 | 23 | 1 | t, r | 2 | NA |
| Morgan et al. (100) Late Puberty | 2 | NA | 2 | 40 | 1 | r | 2 | NA |
| Telzer et al. (104) | 2 | NA | 2 | 39 | 2 | B, β | NA | NA |
| Hasler et al. (93) | 2 | NA | 1 | 93 | 1 | r | NA | NA |

Table S4: Assessment of prediction quality

[1]Cross validation used for orbital loss model

[2]Non-significant unreported effect size, coefficient was pushed to 0 by elastic net, cross validation was used

[3]Used a bootstrap approach to generate confidence intervals, but did not use it for out of sample testing. DMN rated as 2 and G'OC rated as 1, reconciled to 2

[4]Used a CV method for treatment response as a binary, but not for severity

[5]Cross validated, but unclear which steps

[6]Cross validation scheme not specified

| Study | Preregistration | Pregistration repository | Preregistration ID | Shared Code | Shared Data |
|---|---|---|---|---|---|
| Scult, et al. (102) | 2 | | | 2 | 2 |
| Jin et al. (94) High-risk | 2 | | | 2 | 2 |
| Jin et al. (94) Low-risk | 2 | | | 2 | 2 |
| Flores et al. (91) | 2 | | | 2 | 2 |
| Burkhouse et al. (90) CBT | 1 | ClinicalTrials.gov | NCT01903447 | 2 | 2 |
| Burkhouse et al. (90) Setraline | 1 | ClinicalTrials.gov | NCT01903447 | 2 | 2 |
| Bakker et al. (85) | 1 | trialregister.nl | 3662 | 2 | 2 |
| Luo et al. (98) | 2 | | | 2 | 2 |
| Luo et al. (98) | 2 | | | 2 | 2 |
| Barch et al. (86) | 1 | ClinicalTrials.gov | NCT02076425 | 2 | 2 |
| Barch et al. (86) | 1 | ClinicalTrials.gov | NCT02076425 | 2 | 2 |
| Kujawa et al. (95) | 2 | | | 2 | 2 |
| Langenecker et al. (97) | 2 | | | 2 | 2 |
| Bertocci et al. (87) | 2 | | | 2 | 2 |
| Swartz et al. (103) | 2 | | | 2 | 2 |
| Goldstein et al. (106) | 2 | | | 2 | 2 |
| Burani et al. (89) | 2 | | | 2 | 2 |
| Mackin et al. (99) | 2 | | | 2 | 2 |
| Kujawa et al. (96) | 2 | | | 2 | 2 |
| Walsh et al. (105) | 2 | | | 2 | 2 |
| Queirazza et al. (101) | 2 | | | 2 | 2 |
| Greenberg et al. (92) | 1 | PubMed | PMC6100771, PMC5485858 | 2 | 2 |
| Admon et al. (9) | 2 | | | 2 | 2 |
| Stringaris et al. (70) | 2 | | | 2[1] | 2[1] |
| Bress et al. (88) | 2 | | | 2 | 2 |
| Morgan et al. (100) Early Puberty | 2 | | | 2 | 2 |
| Morgan et al. (100) Late Puberty | 2 | | | 2 | 2 |
| Telzer et al. (104) | 2 | | | 2 | 2 |
| Hasler et al. (93) | 2 | | | 2 | 2 |

Table S5: Assessment of adherence to open science practices

[1]DMN and GO'C disagreed about these ratings since information on data access (https://imagen-europe.com/resources/imagen-dataset/) and code (https://github.com/imagen2) is available online, but was not referenced in the publication.

| Modality | Specificity | Design | N | r (95% CI) | z | p | $i^2$ | Worst r | Worst z | Worst p |
|---|---|---|---|---|---|---|---|---|---|---|
| fMRI | Striatum | Treat | 2 | | | | | | | |
| EEG | RewP | Treat | 6 | -0.16 [-0.26, -0.05] | -2.85 | 0.0044 | 20.15% | -0.13 | -1.91 | 0.057 |
| fMRI | Global | Treat | 6 | 0.30 [0.17, 0.42] | 4.37 | | 27.66% | 0.25 | 2.96 | |
| EEG | Global | Treat | 6 | 0.19 [0.09, 0.28] | 3.85 | | 20.89% | 0.17 | 2.88 | |

Table S6: Summary of predictive meta-analytic hypotheses of treatment effects. The “global” results are best-case analyses taking the absolute value of strongest effect from any reward related analysis to define the upper bounds of the relationship between reward processing and future changes in depression. p-values are not given because significant difference from 0 is trivial after taking the absolute value. The least significant results from a leave-one-out analysis are shown in the “worst” columns. No meta-analysis was done on striatal fMRI predicting treatment outcomes because only two studies were found.

| Study | Task | Interval | n | Reliability |
|---|---|---|---|---|
| Chase et al., 2015 | Reward | 7 | 37 | 0.284 |
| Fliessbach et al., (107) | Reward (adapted MID) | 8 | 25 | 0.136 |
| Fliessbach et al., (107) | Reward (box guessing) | 8 | 25 | 0.286 |
| Fliessbach et al., (107) | Reward (number guessing) | 8 | 25 | 0.29 |
| Holiga et al., (108) | MID | 14 | 30 | 0.58 |
| Plitcha et al., (109) | Monetary reward anticipation | 15 | 25 | 0.591 |
| Schlagenhauf, (110) | MID | 28 | 10 | 0.502 |
| Keren et al., (111) | MID | 80 | 18 | 0.801 |
| Elliott et al., (112) | MID | 79 | 20 | 0.45 |

Table S7: Test-retest reliability of fMRI measures of neural reward processing based on information collated in Elliott et al. (6).

| Study | Assessment | Population | r | n | Interval |
|---|---|---|---|---|---|
| Langvik, E. et al. (113) | SHAPS | psychology students | 0.71 | 94 | 70 |
| Watson, D (114) | IDAS-II-Dysphoria | college students | 0.74 | 841 | 14 |
| Sprinkle et al. (115) | BDI-II | undergraduates with initial appointment at a clinic | 0.96 | 46 | 3.2 |
| Harvey, P.D et al. (116) | ALS-anxiety-depression | undergraduate female students | 0.57 | 28 | 28 |
| Harvey, P.D et al. (116) | ALS-anxiety-depression | undergraduate male students | 0.81 | 26 | 28 |
| Gerson et al. (117) | CALS | inpatients of child and adolescent psychiatric... | 0.68 | 35 | 14 |

| | | | | | |
|---|---|---|---|---|---|
| Gerson et al. (117) | CALS | students suburban school | 0.89 | 72 | 14 |
| CF Saylor et al (118) | CDI | school students 5th-6th graders | 0.38 | 69 | 7 |
| CF Saylor et al (118) | CDI | children with emotional problems | 0.87 | 30 | 7 |
| CF Saylor et a.l (118) | CDI | children with emotional problems | 0.59 | 24 | 42 |
| Smucker, M.R et al. (119) | CDI | female elementary school students | 0.74 | 78 | 21 |
| Smucker, M.R et al. (119) | CDI | male elementary school students | 0.77 | 77 | 21 |
| Achenbach TM (120) | YSR(6-18)-internalizing subscale | non referred children | 0.8 | 89 | 8 |

Table S8: Test-retest reliability of clinical symptom measures found by informal review. Interval is the interval between assessments in days.

**# Supplemental Figures**

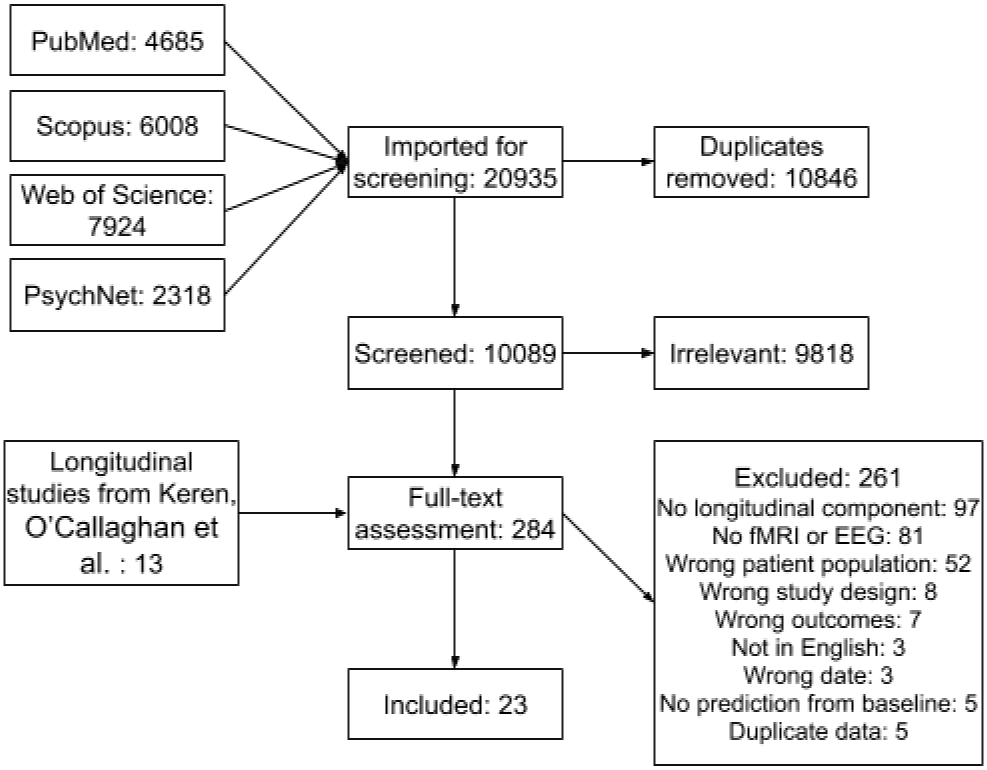


Supplemental Figure S1: PRISMA diagram for review of longitudinal studies of reward processing and depression.

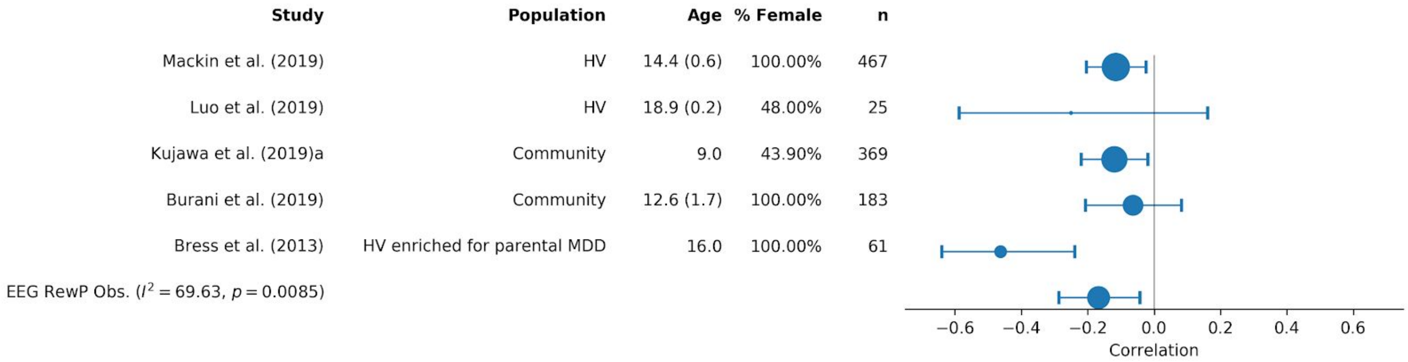


Supplemental Figure S2: Forest plot for random effects meta-analysis of observational EEG studies reporting a reward positivity (RewP) effect for the correlation with change in depressive symptoms.

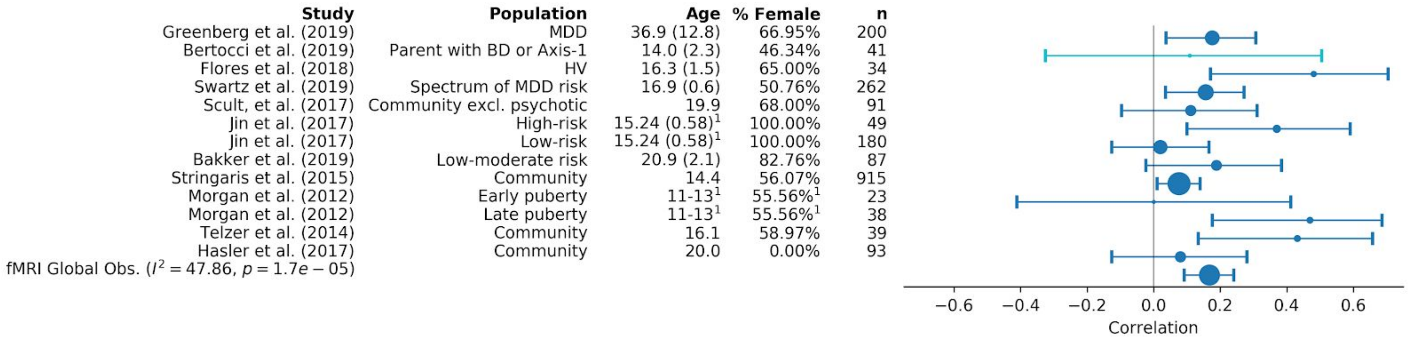


Supplemental Figure S3: Forest plot for random effects meta-analysis of observational fmri studies reporting any effect for the correlation with change in depressive symptoms. Since we were comparing across activation, psychophysiological interactions, and changes in connectivity, we took the absolute value of the reported effects. p-values should be disregarded because significant difference from 0 is trivial after taking the absolute value.

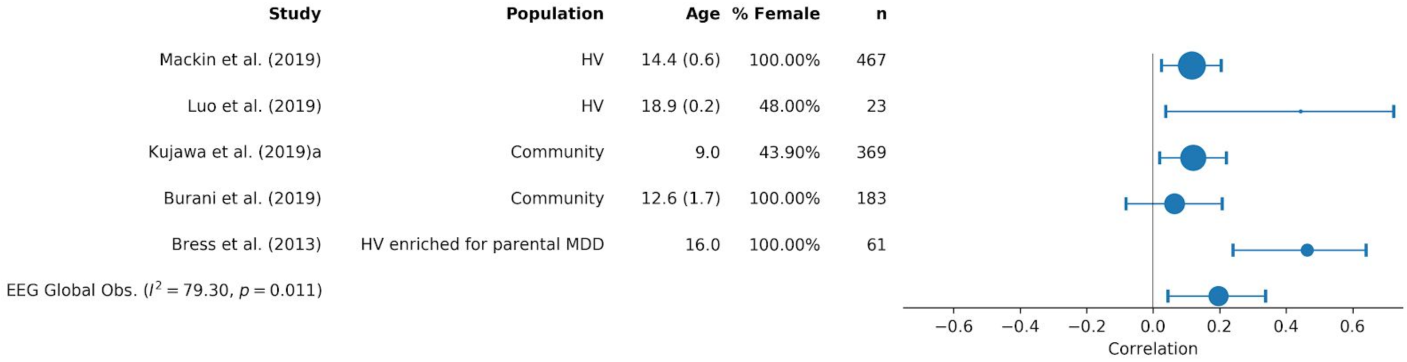


Supplemental Figure S4: Forest plot for random effects meta-analysis of observational EEG studies any effect for the correlation with change in depressive symptoms.  Since we were comparing across multiple signals and analyses, we took the absolute value of the reported effects. p-values should be disregarded because significant difference from 0 is trivial after taking the absolute value.

| Study | Population | Treatment | Age | % Female | n |
|---|---|---|---|---|---|
| Kujawa et al. (2019)a | Community | maternal depression | 9.0 | 43.90% | 369 |
| Kujawa et al. (2019)b | Anxiety disorder + comorbidity | CBT or sertraline | 13.1 (4.0) | 40.70% | 22 |
| Burkhouse et al. (2018) | R-DOC internalizing symptoms | CBT | 28.7 (8.9) | 73.50% | 34 |
| Burkhouse et al. (2018) | R-DOC internalizing symptoms | sertraline | 24.9 (8.1) | 75.90% | 29 |
| Burani et al. (2019) | Community | sleep and stress | 12.6 (1.7) | 100.00% | 183 |
| Barch et al. (2019) | MDD | Parent-Child Interaction Therapy | 5.5 (0.8) | 35.00% | 60 |
| EEG RewP Treat ($I^2 = 19.90$, $p = 0.0041$) | | | | | |

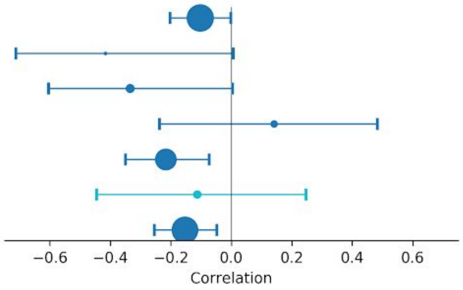


Supplemental Figure S5: Forest plot for random effects meta-analysis of treatment studies with EEG reporting a reward positivity (RewP) effect for the correlation with change in depressive symptoms.

| Study | Population | Treatment | Age | % Female | n |
|---|---|---|---|---|---|
| Greenberg et al. (2019) | MDD | sertraline or placebo | 36.9 (12.8) | 66.95% | 194 |
| Walsh et al. (2017) | MDD | BA | 33.0 (7.1) | 71.05% | 186 |
| Queirazza et al. (2019) | MDD | computerized CBT | 39.2 (12.9) | 48.65% | 26 |
| Bakker et al. (2019) | Low-moderate risk | reward anticipation on activity pleasantness | 20.9 (2.1) | 82.76% | 87 |
| Admon et al. (2015) | MDD | SAMe (5), escitalopram (5), placebo (4) | Adult | 50.00%[1] | 14 |
| Langenecker et al. (2019) | MDD | duloxetine | 28.1 (9.9) | 50.00% | 10 |
| fMRI Global Treat ($I^2 = 28.35$, $p = 9.2e-06$) | | | | | |

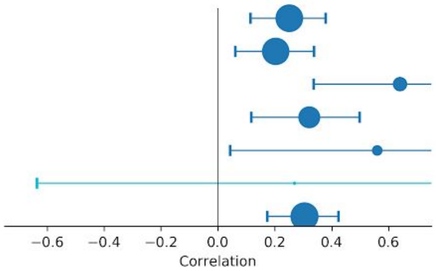


Supplemental Figure S6: Forest plot for random effects meta-analysis of treatment studies with fMRI reporting any effect for the correlation with change in depressive symptoms. Since we were comparing across activation, psychophysiological interactions, and changes in connectivity, we took the absolute value of the reported effects. p-values should be disregarded because significant difference from 0 is trivial after taking the absolute value.

| Study | Population | Treatment | Age | % Female | n |
|---|---|---|---|---|---|
| Kujawa et al. (2019)a | Community | maternal depression | 9.0 | 43.90% | 369 |
| Kujawa et al. (2019)b | Anxiety disorder + comorbidity | CBT or sertraline | 13.1 (4.0) | 40.70% | 22 |
| Burkhouse et al. (2018) | R-DOC internalizing symptoms | CBT | 28.7 (8.9) | 73.50% | 34 |
| Burkhouse et al. (2018) | R-DOC internalizing symptoms | sertraline | 24.9 (8.1) | 75.90% | 29 |
| Burani et al. (2019) | Community | sleep and stress | 12.6 (1.7) | 100.00% | 183 |
| Barch et al. (2019) | MDD | Parent-Child Interaction Therapy | 5.5 (0.8) | 35.00% | 60 |
| EEG Global Treat ($I^2 = 20.89$, $p = 0.00012$) | | | | | |

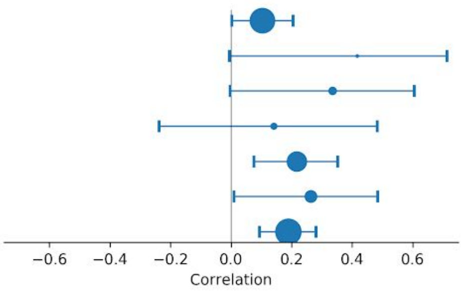


Supplemental Figure S7: Forest plot for random effects meta-analysis of treatment studies with EEG reporting any effect for the correlation with change in depressive symptoms. Since we were comparing across activation, psychophysiological interactions, and changes in connectivity, we took the absolute value of the reported effects. p-values should be disregarded because significant difference from 0 is trivial after taking the absolute value.

Neural correlates of perception of emotional facial expressions in out-patients with mild-to-moderate depression and anxiety. A multicenter fMRI study. *Psychol Med* 41: 2253–2264.

Event-Related Functional Magnetic Resonance ImagingStudy. *Arch Gen Psychiatry* 61: 877–889.

31. Fu CHY, Williams SCR, Brammer MJ, Suckling J, Kim J, Cleare AJ, *et al.* (2007): Neural Responses to Happy Facial Expressions in Major Depression Following Antidepressant Treatment. *Am J Psychiatry* 164: 599–607.
32. Fu CHY, Williams SCR, Cleare AJ, Scott J, Mitterschiffthaler MT, Walsh ND, *et al.* (2008): Neural Responses to Sad Facial Expressions in Major Depression Following Cognitive Behavioral Therapy. *Biol Psychiatry* 64: 505–512.
33. Gorka SM, Huggins AA, Fitzgerald DA, Nelson BD, Phan KL, Shankman SA (2014): Neural response to reward anticipation in those with depression with and without panic disorder. *J Affect Disord* 164: 50–56.
34. Gotlib IH, Sivers H, Gabrieli JDE, Whitfield-Gabrieli S, Goldin P, Minor KL, Canli T (2005): Subgenual anterior cingulate activation to valenced emotional stimuli in major depression. *NeuroReport* 16: 1731–1734.
35. Gotlib IH, Hamilton JP, Cooney RE, Singh MK, Henry ML, Joormann J (2010): Neural Processing of Reward and Loss in Girls at Risk for Major Depression. *Arch Gen Psychiatry* 67: 380–387.
36. Gradin VB, Kumar P, Waiter G, Ahearn T, Stickle C, Milders M, *et al.* (2011): Expected value and prediction error abnormalities in depression and schizophrenia. *Brain* 134: 1751–1764.
37. Gradin VB, Pérez A, MacFarlane JA, Cavin I, Waiter G, Engelmann J, *et al.* (2015): Abnormal brain responses to social fairness in depression: an fMRI study using the Ultimatum Game. *Psychol Med* 45: 1241–1251.
38. Hägele C, Schlagenhauf F, Rapp M, Sterzer P, Beck A, Bermpohl F, *et al.* (2015): Dimensional psychiatry: reward dysfunction and depressive mood across psychiatric disorders. *Psychopharmacology (Berl)* 232: 331–341.
39. Hall GBC, Milne AMB, MacQueen GM (2014): An fMRI study of reward circuitry in patients with minimal or extensive history of major depression. *Eur Arch Psychiatry Clin Neurosci* 264: 187–198.
40. Johnston BA, Tolomeo S, Gradin V, Christmas D, Matthews K, Douglas Steele J (2015): Failure of

hippocampal deactivation during loss events in treatment-resistant depression. *Brain* 138: 2766–2776.

*Psychopharmacol (Oxf)* 26: 1424–1433.

treatment with escitalopram. *J Psychopharmacol (Oxf)* 26: 677–688.

disorder. *Prog Neuropsychopharmacol Biol Psychiatry* 91: 38–48.

98. Luo Y, Jiang H, Chen X, Zhang Y, You X (2019): Temporal dynamics of hedonic and eudaimonic reward processing: An event-related potentials (ERPs) study. *Int J Psychophysiol* 137: 63–71.
99. Mackin DM, Kotov R, Perlman G, Nelson BD, Goldstein BL, Hajcak G, Klein DN (20190502): Reward processing and future life stress: Stress generation pathway to depression. *J Abnorm Psychol* 128: 305.
100. Morgan JK, Olino TM, McMakin DL, Ryan ND, Forbes EE (2013): Neural response to reward as a predictor of increases in depressive symptoms in adolescence. *Neurobiol Dis* 52: 66–74.
101. Queirazza F, Fouragnan E, Steele JD, Cavanagh J, Philiastides MG (2019): Neural correlates of weighted reward prediction error during reinforcement learning classify response to cognitive behavioral therapy in depression. *Sci Adv* 5: eaav4962.
102. Scult MA, Knodt AR, Radtke SR, Brigidi BD, Hariri AR (2019): Prefrontal Executive Control Rescues Risk for Anxiety Associated with High Threat and Low Reward Brain Function. *Cereb Cortex* 29: 70–76.
103. Swartz JR, Weissman DG, Ferrer E, Beard SJ, Fassbender C, Robins RW, *et al.* (2020): Reward-Related Brain Activity Prospectively Predicts Increases in Alcohol Use in Adolescents. *J Am Acad Child Adolesc Psychiatry* 59: 391–400.
104. Telzer EH, Fuligni AJ, Lieberman MD, Galván A (2014): Neural sensitivity to eudaimonic and hedonic rewards differentially predict adolescent depressive symptoms over time. *Proc Natl Acad Sci* 111: 6600–6605.
105. Walsh E, Carl H, Eisenlohr-Moul T, Minkel J, Crowther A, Moore T, *et al.* (2017): Attenuation of Frontostriatal Connectivity During Reward Processing Predicts Response to Psychotherapy in Major Depressive Disorder. *Neuropsychopharmacology* 42: 831–843.
106. Goldstein BL, Kessel EM, Kujawa A, Finsaas MC, Davila J, Hajcak G, Klein DN (undefined/ed): Stressful life events moderate the effect of neural reward responsiveness in childhood on depressive symptoms in adolescence. *Psychol Med* 1–8.

117. Gerson AC, Gerring JP, Freund L, Joshi PT, Capozzoli J, Brady K, Denckla MB (1996): The Children's

Affective Lability Scale: A psychometric evaluation of reliability. *Psychiatry Res* 65: 189–198.

118. Saylor CF, Finch AJ, Spirito A, Bennett B (19850101): The Children's Depression Inventory: A systematic evaluation of psychometric properties. *J Consult Clin Psychol* 52: 955.

119. Smucker MR, Craighead WE, Craighead LW, Green BJ (1986): Normative and reliability data for the children's depression inventory. *J Abnorm Child Psychol* 14: 25–39.

120. Achenbach TM, Rescorla LA (2001): *Manual for the ASEBA School-Age Forms and Profiles*. Burlington, VT: University of Vermont Research Center for Children, Youth, & Families.
